# Chlamydia coinfection inhibits HPV-induced safeguards of the cellular and genomic integrity in patient-derived ectocervical organoids

**DOI:** 10.1101/2021.04.15.439996

**Authors:** Stefanie Koster, Rajendra Kumar Gurumurthy, Hilmar Berger, Marina Drabkina, Hans-Joachim Mollenkopf, Christian Goosmann, Volker Brinkmann, Zachary Nagel, Mandy Mangler, Thomas F Meyer, Cindrilla Chumduri

## Abstract

Cervical mucosa is continually confronted by coinfections with pathogenic microbes. In addition to human papillomavirus, coinfections with *Chlamydia trachomatis* have been associated with an increased risk of cervical cancer. However, the dynamics of coinfections, their impact on the epithelia, and their contribution to pathogenesis remain obscure. Using a novel human ectocervical squamous stratified epithelial organoids, we recapitulated the natural infections of the cervix by Chlamydia, HPV, and their coinfections. Towards this, we genetically manipulated the healthy organoids to mimic in vivo HPV persistence by introducing E6E7 oncogenes into the host genome. HPV persistent organoids show enhanced tissue regeneration, increased proliferation and differentiation of stem cells, and nuclear atypia resembling cervical intraepithelial neoplasia grade 1. We found that HPV interferes with normal Chlamydia development. Further, a unique transcriptional host response induced by Chlamydia and HPV leads to distinct reprogramming of host cell processes. Strikingly, in coinfections, Chlamydia impedes HPV-induced mechanisms that maintain cellular and genome integrity, including mismatch repair (MMR). Distinct post-translational proteasomal-degradation and E2F-mediated transcriptional regulation delineate the inverse regulation of MMR during coinfections. Our study employing organoids demonstrates the jeopardy of multiple sequential infection processes and the unique cellular microenvironment they create, accelerating neoplastic progression.

## Introduction

Cervical mucosa being a protective barrier against invading pathogens into the upper female reproductive tract is often challenged by the invading pathogens and dysbiosis (Kwasniewski, Wolun-Cholewa et al. 2018, Chumduri and Turco 2021). Coinfections are increasingly being discovered in clinical practice, yet their implication is not always clear. The lack of in vitro systems that recapitulate the human cervical epithelium has been a bottleneck to decipher the individual and coinfection processes systematically and elucidate their role and impact in driving pathogenesis, including cervicitis and carcinogenesis. The recently established human and mouse-derived primary cervical epithelial three-dimensional (3D) organoids mimic the in vivo native tissue architecture and can be expanded and maintained long term. Thus, they offer a unique possibility to elucidate the dynamics and the impact of different infections and coinfections in cervical pathogenesis (Chumduri, Gurumurthy et al. 2018, Chumduri, Gurumurthy et al. 2021, Chumduri and Turco 2021). The cervix comprises ectocervix lined by stratified squamous epithelium that projects into the vagina, and the endocervix lined by columnar epithelium that forms a continuum with the uterus. The ecto- and endocervix merge at the squamocolumnar transition zone and these sites are highly predisposed to preneoplastic squamous metaplasia, a precursor of cervical cancers (Kwasniewski, Wolun-Cholewa et al. 2018, Chumduri, Gurumurthy et al. 2021, Chumduri and Turco 2021). Most cervical cancers, the fourth most common cancers of women worldwide, originate from the squamous stratified epithelium (Burghardt and Ostor 1983, Chumduri, Gurumurthy et al. 2021).

Human papillomavirus (HPV) and bacterial pathogen *Chlamydia trachomatis* (*C. trachomatis*) are among the highly prevalent sexually transmitted infections. HPV has long been established as the etiological agent of cervical carcinogenesis (Durst, Gissmann et al. 1983, zur Hausen 2009). Although HPV infections are encountered by over 80 percent of women during their lifetime, less than two percent with HPV infections develop cervical cancer (Burchell, Winer et al. 2006, Dunne, Unger et al. 2007). Cofactors like immune status, hormones, and coinfections are emerging as a causal link in cervical cancer development (Koskela, Anttila et al. 2000, Brake and Lambert 2005, Zhu, Shen et al. 2016). Coinfections with *C. trachomatis* are seen at an increased incidence in patients with invasive cervical and ovarian cancers (Koskela, Anttila et al. 2000, Ssedyabane, Amnia et al. 2019), yet the coinfection dynamics and the underlying mechanisms are entirely unknown. Unlike tumor viruses whose DNA can be found within the tumors, bacteria linked to cancers rarely leave any traceable elements. The HPV persistence by integrating oncogenes E6E7 into the host genome is a major trigger and point of no return during cervical cancer development. One link to associate bacteria with the onset of cancers is identifying the cellular and mutational processes that contribute to cell transformation.

Previous studies performed in immortalized or cancer cell lines, although they may not completely reflect near-physiological processes, led to major advances in understanding the impact of HPV E6E7 oncogenes or Chlamydia on the infected cells. HPV E6E7 oncogenes regulate several host processes that counteract anti-proliferative signals by inhibiting the cell cycle arrest to support viral DNA replication (Beglin, Melar-New et al. 2009, Moody and Laimins 2010, Zhong, Bechill et al. 2015). The HPV E7 mediates E2F transcriptional factor activation by releasing it from inhibitory interaction with Retinoblastoma (RB) protein, while E6 promotes tumor suppressor TP53 protein degradation, thus avoiding growth arrest and promote proliferation (Scheffner, Werness et al. 1990, Scheffner, Munger et al. 1992, Bracken, Ciro et al. 2004, Poppy Roworth, Ghari et al. 2015). *C. trachomatis* promotes cell proliferation by activating MAPK, PI3K-AKT, and blocking apoptosis (Gonzalez, Rother et al. 2014, Chumduri, Gurumurthy et al. 2016). It enhances epithelial to mesenchymal transition by controlling the host transcription factors downstream of MAPK, fostering migratory advantage to the infected cells (Zadora, Chumduri et al. 2019). *C. trachomatis* induces DNA doublestrand breaks and 8-oxo-dG lesions. However, it inhibits the genome surveillance, including cell cycle checkpoint and high-fidelity homologous recombination repair by modulating the host protein phosphatases and suppressing the activation of ATM signaling (Chumduri, Gurumurthy et al. 2013, Chumduri, Gurumurthy et al. 2016, Mi, Gurumurthy et al. 2018). HPV, unlike *C. trachomatis*, activates ATM and ATR signaling and recruits DNA repair factors, which are essential for viral replication (Moody and Laimins 2009, Anacker and Moody 2017, Ngoi, Sundararajan et al. 2020). However, HPV uncouples downstream cell cycle checkpoint responses by modulating cyclins and cyclin-dependent kinase (CDK) inhibitors to maintain cell proliferation (Spardy, Covella et al. 2009, Moody and Laimins 2010). These studies suggest that each pathogen reprograms host cellular processes distinctively to facilitate their propagation within the host cell. However, the coinfections with HPV and *C. trachomatis* might elicit more complex host cell responses.

This study addressed two key aspects; first, we introduce a near-physiological human ectocervical organoid model to study the interaction of stratified epithelial tissue barrier and infections. Second, we modeled individual and coinfection dynamics of persistent HPV and Chlamydia and their impact on the host processes. Human ectocervical organoids derived from HPV-negative healthy donors were genetically manipulated to integrate HPV16 E6E7 DNA into the stem cell genome. These HPV negative and HPV E6E7 expressing organoids were infected with *C. trachomatis* and characterized the infection process in a single and coinfection scenario. HPV E6E7 integration induces low-grade neoplastic changes resembling cervical intraepithelial neoplasia grade 1 (CIN1) and promotes aberrant chlamydia development. The global transcriptomic analysis revealed HPV and *C. trachomatis-induced* unique host cellular reprogramming. Several genes were discovered to be similarly up or down-regulated by both pathogens involving specific immune responses. Strikingly, a significant subset of all regulated genes controlled by an E2F transcription factor and associated with DNA mismatch repair (MMR) was discovered to be oppositely regulated by HPV and *C. trachomatis* at the transcriptional and post-translational level.

Interestingly, two of the seven identified mutational signatures in cervical cancers (COSMIC, Tate, Bamford et al. 2019) are attributed to defective MMR. Improper DNA repair in stem cells can lead to the formation and accumulation of somatic mutations that can subsequently be transferred to the daughter cells and are linked to cancer development (Weeden and Asselin-Labat 2018). This study goes beyond the state-of-the-art and shows the hazards posed by multiple sequential infections and the molecular mechanisms contributing to carcinogenesis. Our cervical organoids prove to be excellent in vitro model for studying complex interactions between epithelial tissue and pathogens and analyzing the molecular sequels of these initial events of pathogenesis.

## Results

### Human ectocervical three-dimension organoids to mimic HPV persistence

To overcome the current lack of a suitable epithelial primary cell model that recapitulates the human ectocervix’s stratified epithelium and model disease development, we recently established an adult stem cell-derived-ectocervical organoids (Chumduri, Gurumurthy et al. 2018, Chumduri, Gurumurthy et al. 2021). The adult ectocervical stem cells isolated from healthy donors were embedded in Matrigel to culture 3D organoids. Alternatively, they were first cultured in collagen-coated cell culture flask with a defined cocktail of growth factors to enrich stem cells and were subsequently cultured on irradiated mouse fibroblast (J2-3T3) to maintain the stem cells in long-term cultures (Figure 1A). These stem cells were used for experiments or to generate the mature ectocervical 3D organoids comprising stem cells and differentiated cells.

**Figure 1:**
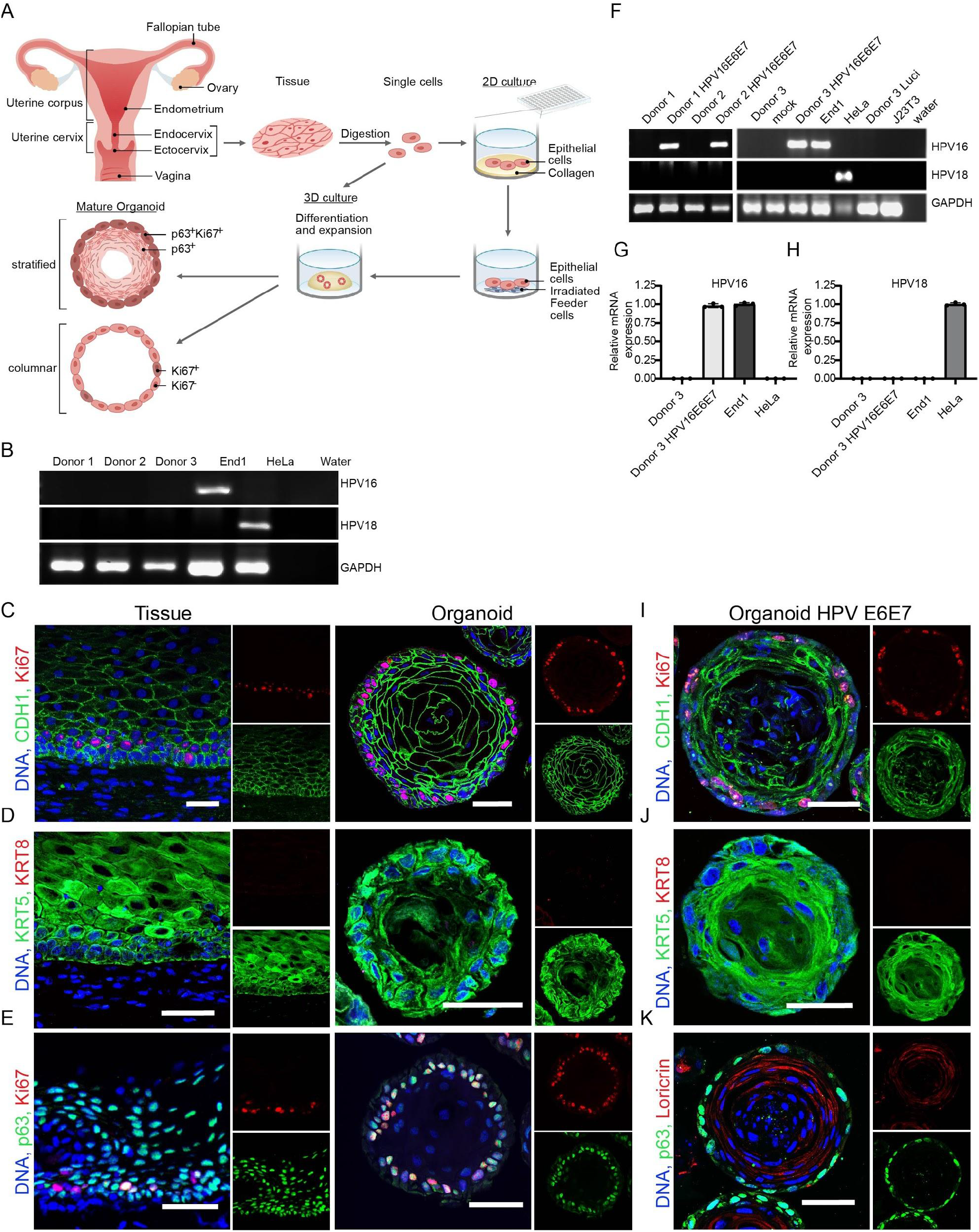
Genetic manipulation of HPV negative ectocervical organoids to mimic persistent HPV infection. (A) Schematic representation of human ectocervical organoid and 2D stem culture generation from biopsies. (B) Ectocervical organoids from three healthy donors were subjected to a polymerase chain reaction (PCR) to detect HPV 16 and HPV 18 E6E7. HeLa and End 1 cells were used as positive controls for HPV 18 and HPV 16, respectively. (C-E) Representative confocal images of human ectocervical tissue and organoids immunolabeled for the marker of proliferation Ki67, epithelial E-cadherin (CDH1) (C); squamous lineage KRT5, columnar lineage KRT8 (D); Ki67, basal stem cell p63 (E), Nuclei (Hoechst) are shown in blue. (F) PCR to detect HPV16, HPV18 E6E7, and GAPDH. Donor 1-3, untreated in human ectocervical stem cells; mock, cells treated with Polybrene; Donor 1-3 HPV E6E7, cells from Donor 1-3 transduced with lentivirus carrying HPV16 E6E7; End1, End1 cells; HeLa, HeLa cells; Donor 3 Luci, cells transduced with lentivirus carrying Luciferase; J23T3, feeder cells used for stem cell culture; water, water control. (G-H) Relative mRNA expression of HPV16 E6E7 (G) and HPV18 E6E7 (H) genes analyzed by quantitative real-time PCR from Donor 3. Data represent means ± standard deviations (SD) of results normalized to positive controls End1 and HeLa cells, respectively. (I-K) Representative confocal images of human ectocervical organoids expressing HPV16 E6E7 immunolabeled for Ki67, CDH1 (I); KRT5, KRT8 (J); p63, differentiation marker Loricrin (K), Nuclei (Hoechst) are shown in blue. Images in C-E and I-K represent organoid cultures from n = 3 biologically independent experiments. Scale bar= 50 μm.

To systematically evaluate the impact of persistent HPV infection and subsequent coinfection with *C. trachomatis* on the host regulation, it is essential to have epithelial organoids from individuals negative for high-risk HPV (hrHPV) integrations. Therefore, we established ectocervical organoids from healthy donors and tested for E6E7 genes from various hrHPV types. Shown are the ectocervical organoids from three donors found to be negative for E6E7 oncogenes of all tested hrHPV, including HPV 16 and 18 (Figure 1B), which account for over 70% of cervical cancers (Shanmugasundaram and You 2017). The healthy human ectocervical (hCEcto) tissue-derived lineage-specific cytokeratin 5 positive (KRT5+) epithelial stem cells can self-organize into 3D organoids resembling the stratified squamous epithelium of the parent tissue. These organoids comprise the basal progenitor, parabasal, intermediate, and non-keratinized terminally differentiated superficial layers. They express epithelial marker E-cadherin (CDH1) and are KRT5+ but negative for KRT8, a columnar epithelium marker of the endocervix. The outer layer of the organoid comprises p63+ and Ki67+ basal progenitors. They have the proliferative and regenerative capacity to produce non-keratinized differentiated epithelium by the continuous movement of cells from the basal to luminal layers (Figure 1C-E) (Chumduri, Gurumurthy et al. 2018, Chumduri, Gurumurthy et al. 2021).

Next, the organoids from these healthy donors without E6E7 from hrHPV strains were used to study the impact of HPV integration on the healthy cervical tissue. Towards this end, the hCEcto stem cells were transduced with lentiviruses carrying HPV16 E6E7 oncogenes to genetically modify these cells to integrate E6E7 into the host cell genome. As expected, the untransduced hCEcto stem cells from donors 1-3 were negative for both HPV16 and HPV18, while the hCEcto cells transduced with lentiviruses were positive for HPV16 E6E7 in all three donors (Figure 1F-H).

These hCEcto cells expressing HPV16 E6E7 (hCEcto E6E7) preserve their stemness and retain their capacity to grow into mature organoids under the same culture conditions as hCEcto epithelial stem cells. The hCEcto E6E7 cells could be expanded and long-term propagated both as 2D stem cell cultures on the feeder system and as fully matured organoids. The two-week-old hCEcto E6E7 mature organoids consist of an outer layer of basal p63+ progenitors that are proliferative (Ki67+) and remained KRT5+/KRT8- (Figure 1I-K) similar to the normal ectocervical epithelium. However, a significant alteration in the distribution of adherens junction marker E-cadherin (CDH1), responsible for maintaining cell polarity and differentiation, was observed in E6E7 positive organoids. E-cadherin expression in these organoids was brightest in the basal and parabasal layers, unlike in E6E7 negative organoids (Figure 1I). hCEcto E6E7 organoids were found to proliferate faster, producing several layers of differentiated cells with a striking change from non-keratinized loricrin negative differentiated cells, a phenotype of normal ectocervical epithelial tissue (Dinh, Okocha et al. 2012), to increased keratinization of the differentiated cells, as shown by the presence of loricrin (Figure 1K). They also show nuclear atypia with varied nuclear size (Figure 1I-K). Together, integrating E6E7 into ectocervical organoids induces a phenotype with features reminiscent of cervical intraepithelial neoplasia grade 1 (Robert J. Kurman 2011, Liu, Qian et al. 2015).

### Modelling *C. trachomatis* coinfection in HPV persistent organoids

Next, we sought to establish the ectocervical organoids as *C. trachomatis* infection model and investigate if these organoids support *C. trachomatis* infection and developmental life cycle. For this purpose, we first established *C. trachomatis* infection protocols in the ectocervical organoids. *C. trachomatis* has a biphasic life cycle. The non-replicative elementary bodies (EBs) infect the cells and transform to non-infectious replicative reticulate bodies (RBs). At the time of exit from the host cells, these RBs redifferentiate to EBs to initiate new infections. Efficient infection of both hCEcto and hCEcto E6E7 organoids was achieved by incubating intact small organoids (5 days old) for 2 hours with *C. trachomatis* EBs and reseeding them back into Matrigel to allow these organoids to continue their growth. To follow the progression of *C. trachomatis* infection in organoids, we used recombinant *C. trachomatis* L2 strain expressing enhanced green fluorescent protein (GFP-*Ctr L2*) for ease of detection in the organoids (Figure 2A). First, GFP signals from the developing inclusion were detected 24 hours post-infection (hpi) in basal and parabasal cells. At 5 days post-infection (dpi), strong GFP signals were detected across all differentiated cell layers, and inclusion size increased significantly (Figure 2A; Supplementary Figure S1A). The progression of *C. trachomatis* infection from the basal stem cell layer was observed to be bidirectional. It progressed from the basal cells to daughter cells that self-renew by symmetric division and remain in the basal layer as stem cells, and the ones that form the differentiated epithelial cells via asymmetric division of the stem cells and move towards the lumen. Thus, infected stem cells can be a potential reservoir for *C. trachomatis* to propagate infection in the host. Further, we performed an infectivity assay to evaluate if *C. trachomatis* can complete its developmental cycle by redifferentiation from RBs to EBs within the organoids. The EBs obtained from the lysate of infected organoids at 1dpi or 5dpi were used to infect a monolayer of HeLa cells. Quantification of *C. trachomatis* inclusion forming units revealed a ten-fold increase in infectivity from lysates-derived from organoids infected for 5 days compared to 1 day (Figure 2B), reflecting the increase in the infection load with time. A significant increase in inclusion size and infectious progenies shows *C. trachomatis* replicative activity and completion of its developmental life cycle in ectocervical organoids.

**Figure 2:**
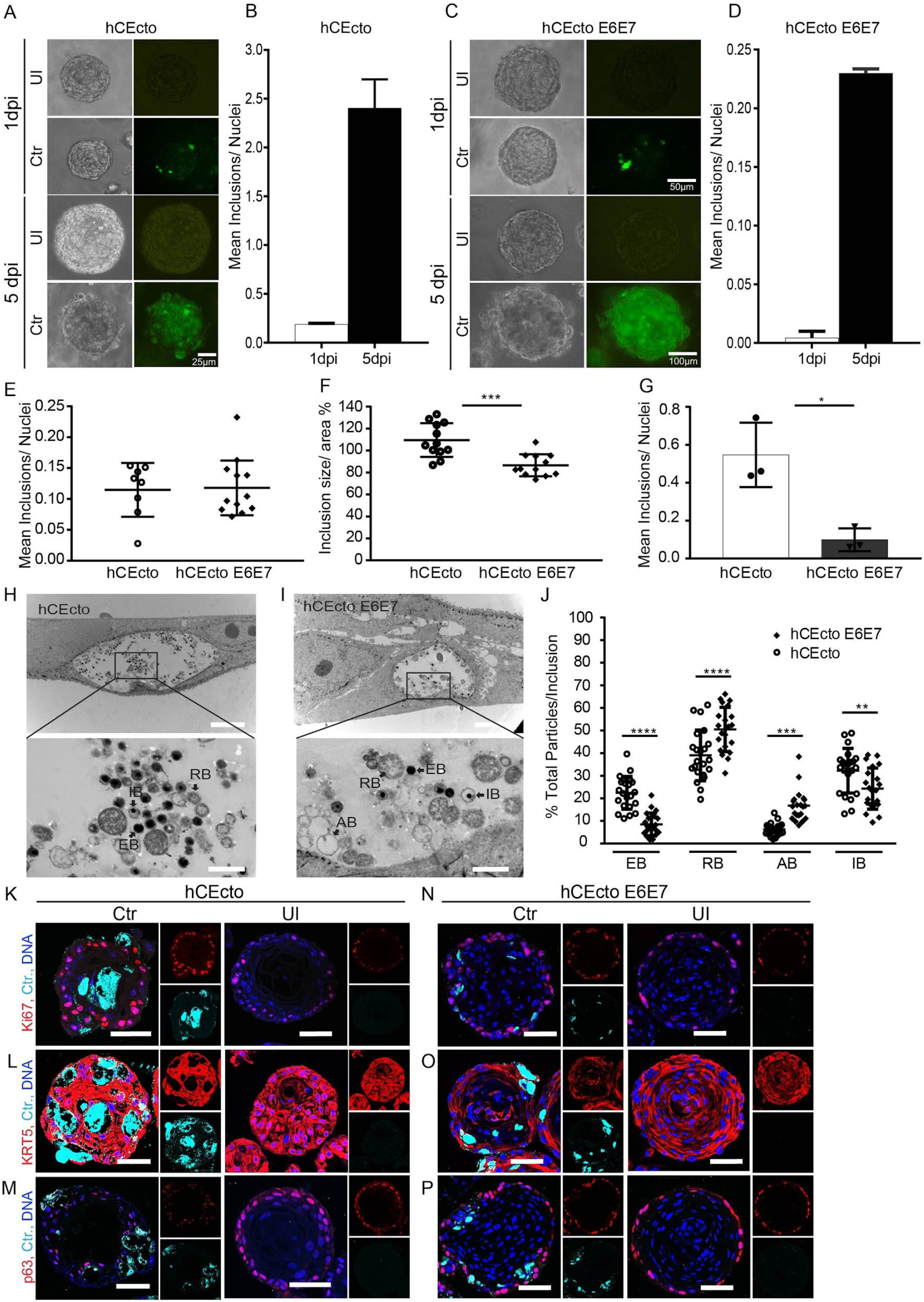
Modelling *C. trachomatis*, HPV16 persistence, and coinfection in human ectocervical organoids. (A) Representative bright-field and fluorescent images of healthy ectocervical (hCEcto) organoids at 1 and 5 dpi with GFP expressing *C.trachomatis* (Ctr). (B) Quantification of infectivity of the lysates derived from Ctr infected hCEcto organoids at 1 and 5 dpi. Shown is the mean ± SD of two replicates representative of three biological replicates. (C) Representative bright-field and fluorescent images of ectocervical organoids with E6E7 expression (hCEcto E6E7) at 1 and 5 dpi with GFP expressing Ctr. (D) Quantification of infectivity of the lysates derived from Ctr infected hCEcto E6E7 organoids at 1 and 5 dpi. Shown is the mean ± SD of two replicates representative of three biological replicates. Images in A and C represent three biological replicates. (E-F) Primary infection of Ctr infected hCEcto and hCEcto E6E7 cells (E) and relative inclusions size (F). Shown is the mean inclusion size per replicate and ± SD from ≥ 8 technical replicates. (G) Quantification of infectivity of the lysate derived from Ctr infected hCEcto and hCEcto E6E7 2D stem cells at 48 hpi. Shown is the mean ± SD of 3 technical replicates (*p<0.05). (H-J) Transmission electron micrographs of Ctr infected hCEcto (H) and hCEcto E6E7 (I) 2D stem cells at 24hpi. Inclusion contains different development forms of Ctr (RB, reticulate bodies; EB, elementary bodies; IB, intermediate stage; AB, aberrant bodies). Shown are representative images from three biological replicates. Scale bar 5 μm, 1μm (inset). (J) Percentage of total EBs, RBs, IBs, and ABs quantified from over 21 inclusions, representative of three independent experiments (**p<0.01, ***p<0.001, **** p<0.0001). (K-P) Confocal images of hCEcto (K-M) and hCEcto E6E7 (N-P) organoids at 5 dp Ctr infection immunolabeled for Ki67 (K, N); KRT5 (L, O), p63 (M, P) and Ctr-MOMP, Nuclei (Hoechst) are shown in blue. Scale bar= 50 μm.

Next, we investigated the impact of persistent HPV on *C. trachomatis* development. Similar to hCEcto, organoids with HPV persistence showed first detectable inclusions in basal and parabasal cells 1dpi. At 5dpi, inclusion size increased significantly and was detected in all epithelial layers (Figure 2C) with an increase in infection load with time (Figure 2D). Although both hCEcto and hCEcto E6E7 cells could be infected with *C. trachomatis* similarly in organoids or 2D stem cells as shown by inclusion formation (Figure 2E, Supplementary Figure S1C), HPV E6E7 suppressed the intracellular development of *C. trachomatis* shown by smaller inclusion size (Figure 2F, Supplementary Figure S1B). Further, *C. trachomatis* from the HPV E6E7 expressing organoids revealed the loss of reinfection ability compared to hCEcto organoids (Figure 2G; Supplementary Figure S1C), showing that the expression of HPV E6E7 interferes with *C. trachomatis* normal development.

To evaluate the influence of HPV persistence on different development stages of *C. trachomatis*, we performed transmission electron microscopy of Chlamydia infected hCEcto and hCEcto E6E7 cells. The ultrastructural analysis revealed that the inclusions contain a mixture of particles from different *C. trachomatis* developmental stages. In hCEcto cells, reproductive RBs mostly located at the periphery, condensed infective EBs, and the intermediate stage (IB) were detected (Figure 2H). These observations are in line with earlier reports (Swanson, Eschenbach et al. 1975, Phillips, Swenson et al. 1984, Peterson and de la Maza 1988). In hCEcto E6E7, besides the EB, RB, and IBs, we found an increased number of aberrant chlamydial developmental forms (AB). As described previously (Wyrick 2010), these ABs are membrane-lined, roundish, pleomorphic or irregularly shaped, and less electron-dense or hollow because of loss of cytoplasmic content and appear like ghost bacterial structures (Figure 2I). Quantification of the various developmental forms revealed that the proportion of all *C. trachomatis* stages is significantly different between E6E7 positive and negative cells. The number of infectious EBs is significantly reduced in E6E7 expressing primary cells, while the proportion of RBs and ABs is enhanced (Figure 2J), corroborating with the reduced infectivity observed in these cells (Figure 2G). These data reveal that persistent expression of HPV16 E6E7 increases AB formation and simultaneously inhibits RB to EB redifferentiation, resulting in a slowdown of *C. trachomatis* development life cycle initiating persistence.

To assess the effects of *C. trachomatis* on host cell proliferation and epithelial architecture, we performed immunofluorescent staining for Ki67, KRT8, KRT5, p63, and *C. trachomatis* major outer membrane protein (MOMP) in hCEcto and hCEcto E6E7 organoids infected for 5 days with *C. trachomatis.* MOMP staining revealed large inclusions across different epithelial layers and inside the lumen (Figure 2K-P). Both hCEcto and hCEcto E6E7 remain KRT5+ and KRT8- (Figure 2l-O). However, an increased number of Ki67+ cells were observed, indicating an enhanced proliferation in *C. trachomatis* infected organoids compared to uninfected organoids (Figure 2K). A similar pattern of changes was observed in hCEcto E6E7 organoids coinfected with *C. trachomatis* (Figure 2N-P). These results show that *C. trachomatis* infection induces proliferative signals in single and coinfections of HPV E6E7 expressing ectocervical epithelium.

### *C. trachomatis*, HPV E6E7, and coinfection elicit unique transcriptional responses in human ectocervical organoids

Next, we performed the transcriptomic analysis to identify the impact of *C. trachomatis*, HPV E6E7, and coinfection on cellular signaling. Towards this, hCEcto and hCEcto E6E7 cells were infected with *C. trachomatis* for 48h in 2D stem cell cultures or 5 days in the 3D organoids. All genes differentially up or down-regulated (log2FC>2, and adj. p-value < 0.05) upon *C. trachomatis*, HPV E6E7, and coinfection compared to uninfected HPV negative healthy cells were scored. Overall, our analysis revealed a massive transcriptional response to HPV E6E7 expression and *C. trachomatis* infection, respectively (Figure 3 A; Supplementary Figure S2A, Supplementary Table S1). In 2D cultured stem cells, HPV E6E7 differentially regulated 3,456 transcripts, while *C. trachomatis* infection led to differential expression of 7,239 transcripts. Among these, 717 genes are similarly regulated by both pathogens, while 706 genes are oppositely regulated (Figure 3B). In the 3D organoid model, we identified 3,139 transcripts were differentially regulated by HPV E6E7 expression, while *C. trachomatis* infection led to differential expression of 2,742 transcripts; of these, 610 genes are similarly regulated, and 203 genes are oppositely regulated by HPV and *C. trachomatis* (Supplementary Figure S2B). Further, coinfections of HPV E6E7 expressing hCEcto cells with *C. trachomatis* exhibited unique gene expression changes. Strikingly, of the genes upregulated by HPV E6E7, 28% reverted to expression levels similar to that of control or were strongly downregulated when coinfected with *C. trachomatis.*

**Figure 3:**
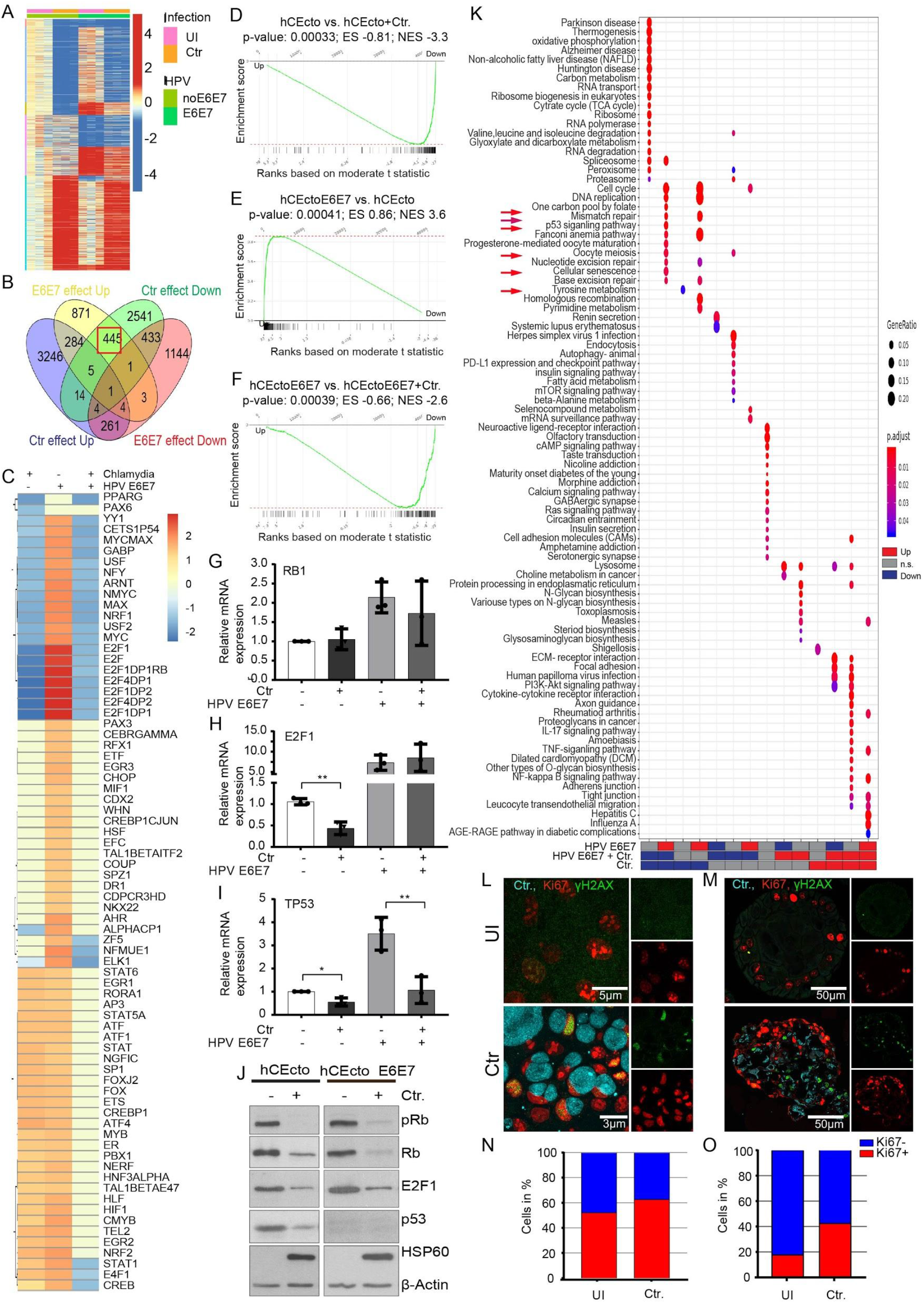
Host transcriptional response in human ectocervical stem cells of *C. trachomatis* and HPV16 E6E7 and coinfection. (A) Heatmap of differentially regulated genes in hCEcto and hCEcto E6E7 2D stems cells with or without Ctr infection with p-value< 0.05; log2 fold change (FC) >1; log2 FC<-1 between any of the conditions from three replicates. (B) Heatmap showing GSEA enrichment (-log10(p-value)) of genes that share Cis-regulatory motifs for transcription factors (stated in the right column). Significant enrichment of genes regulated by E2F1 transcription factor are upregulated by HPV16 E6E7 expression while being downregulated by single Ctr infection and in coinfection. (C) Venn diagram comparing genes significantly (log2 FC ≥ 2, p-value<0.05) up or down-regulated after Ctr infection or HPV E6E7 expression in hCEcto 2D stem cells. Highlighted in the red box is a subset of genes oppositely regulated by both pathogens, upregulated by HPV E6E7 and downregulated by Ctr. (D-F) GSEA analysis on differential gene expression comparisons of hCEcto vs. hCEcto+Ctr, hCEcto vs. hCEcto E6E7, hCEcto E6E7 vs. hCEcto E6E7+Ctr 2D stem cells identified E2F transcription factor target gene to be significantly regulated; downregulated after Ctr infection (D), upregulated upon HPV E6E7 expression (E), and downregulation upon coinfection of HPV E6E7 expressing cells with Ctr (F). (G-J) hCEcto and hCEcto E6E7 2D stem cells were infected with Ctr for 48h. Shown is the relative mRNA expression of RB1 (G), E2F1 (H), and TP53 (I) analyzed by qRT-PCR. Data represent mean ± SD from three biological replicates normalized to uninfected controls. **p<0.01, *p<0.05, Student’s t-test. (J) hCEcto and hCEcto E6E7 2D stem cells were infected with Ctr for 48h, and cell lysates were subjected to immunoblot analysis for indicated proteins and chlamydia HSP60 and β-actin as a loading control. Data represent three biological replicates. (K) Dot plot showing significantly up or down-regulated KEGG pathways among the differentially expressed genes from hCEcto and hCEcto E6E7 2D stem cells with or without Ctr infection categorized into distinct groups as shown. Red arrows highlight DNA repair pathways. Dot diameter refers to the gene ratio within a group. Fill color depicts the adjusted p-value. (L-M) Representative confocal images of hCEcto 2D stem cells (L) and organoids (M) infected for 36 h and 5 d, respectively with Ctr and immunolabelled for the proliferation marker Ki67 (red), DNA damage indicator γH2AX (green), and Chlamydia MOMP (blue). (N-O) Quantification of the number of Ki67 positive cells from confocal images of uninfected and Ctr infected hCEcto 2D stem cells at 48 hpi and (N) organoids at 5 dpi (O).

GSEA analysis revealed several transcription factors (TF) are oppositely regulated by *C. trachomatis* and HPV E6E7. Strikingly, E2F family members were the most significant oppositely regulated TF among all comparisons (Figure 3C; Supplementary Figure S2C, Supplementary Table S2). E2F target genes are significantly upregulated by HPV E6E7 but downregulated by *C. trachomatis* in hCEcto stem cells (Figure 3D-E) and organoids (Supplementary Figure S2D-E). In coinfections, *C. trachomatis* overrides the HPV E6E7 induced E2F target genes expression (Figure 3F, Supplementary Figure S2F). It is known that HPV E7 unleashes the E2F TF from RB inhibitory interaction to activate transcription of downstream targets. However, to date, not much is known regarding Rb-E2F1 pathway modulation by *C. trachomatis.* Further, it is unclear what consequences this *C. trachomatis* mediated suppression of HPV E6E7 induced E2F targets will have on host cells and pathogens.

To investigate the regulation of the Rb-E2F1 pathway by both pathogens, we infected hCEcto and hCEcto E6E7 with *C. trachomatis* and performed quantitative reverse transcription PCR (qRT-PCR) and immunoblot analysis (Figure 3G-J). qRT-PCR analysis revealed that *E2F1* and *TP53* are significantly suppressed by *C. trachomatis* in the absence of HPV E6E7 expression, while coinfection results in a significant reduction of *TP53*, but not *E2F1* and *RB1* transcripts. In contrast to *C. trachomatis* infection, HPV E6E7 upregulated *TP53, E2F1*, and *RB1* gene expression (Figure 3G-I). Immunoblot analysis revealed the reduction of total Rb, pRb (S807/811), E2F1, and p53 proteins upon *C. trachomatis* infection in human ectocervical epithelial cells irrespective of HPV status (Figure 3J). However, E6E7 expression increases E2F1 protein levels and induces near complete degradation of p53 while having no significant effect on total Rb, pRb (S807/811) protein levels. This is consistent with the literature that p53 protein is degraded by HPV E6 (Yim and Park 2005). Further, it is also known that *C. trachomatis* induces degradation of p53 via the PI3K-Akt pathway and MDM2 activation (Gonzalez, Rother et al. 2014, Siegl, Prusty et al. 2014). Together, our data suggest HPV induces E2F1 activation, but Chlamydia suppresses it. HPV and Chlamydia lead to p53 protein degradation. However, at the transcriptional level, p53 is upregulated by HPV while downregulated by Chlamydia in both single and coinfections.

Next, we assessed the biological functions affected by the differentially expressed genes by performing GSEA analysis between uninfected, *C. trachomatis* infected, E6E7 expressing, and coinfected hCEcto cells to identify overrepresented networks from Kyoto Encyclopedia of Genes and Genomes (KEGG) and Gene Ontology (GO) terms. The tumor necrosis factor (TNF) pathway was strongly upregulated by both HPV E6E7 and *C. trachomatis.* Further, specific pathways were uniquely regulated by each pathogen. The inflammatory response (e.g., IL-17 and NF-kappa B signaling) and mitogen-activated protein kinase (MAPK) pathways are distinctly upregulated by *C. trachomatis*, while pathways like oxidative phosphorylation, RNA regulation, and RNA processing are downregulated. The enriched GO terms highlight the opposite regulation of differentiation processes (e.g., skin development and keratinocyte differentiation) and extracellular matrix organization by *C. trachomatis* infection and HPV E6E7. HPV E6E7 expression showed to have a negative impact on differentiation processes, increased mitotic activities, RNA and DNA replication (Supplementary Figure S2H-I).

Intriguingly, pathways involved in DNA replication, DNA damage repair pathways including base excision repair (BER), mismatch repair (MMR), nucleotide excision repair (NER), Fanconi anemia (FA), homologous recombination (HR), and p53 signaling pathway are oppositely regulated by the two pathogens: HPV E6E7 expression activates these pathways, while *C. trachomatis* suppress them (Figure 3K, Supplementary Figure S2G). *C. trachomatis* infection was previously shown to induce DNA double-strand breaks (DSB) in immortalized cells and cancer cell lines (Chumduri, Gurumurthy et al. 2013, Mi, Gurumurthy et al. 2018). Here we show that Chlamydia induces DNA DSB in long-lived stem cells and differentiated cells while promoting cellular proliferation as shown by an increased number of Ki67 and γH2AX positive cells (Figure 3L-O).

### *C. trachomatis* suppresses HPV-induced mismatch repair pathway in ectocervical stem cells

Since the E2F family of transcription factors, vital regulators of several genes involved in cell proliferation and growth arrest, are downregulated by *C. trachomatis* and coinfection, we further analyzed the significance of their altered regulation on the Chlamydia infected cells. RB-like, E2F4, multi-vulval class B proteins and dimerization partner (DP1) form the DREAM complex that mediates gene repression promoting quiescence. Disruption in the DREAM complex-mediated regulation switches the balance from quiescence to proliferation by loss of cell cycle checkpoint gene expression, which is frequently observed in cancers (Sadasivam and DeCaprio 2013, Uxa, Bernhart et al. 2019). Our analysis revealed that many DREAM complex target genes are downregulated by Chlamydia and in coinfection, that could prevent quiescence and promote cell proliferation, which is corroborated by increased Ki67+ cells upon Chlamydia infections (Figure 4A, Figure 3L-O, Supplementary Table 3). Notably, genes involved in DNA repair pathways are among the E2F1 target genes oppositely regulated by Chlamydia and HPV (Figure 4A-B red boxes).

**Figure 4:**
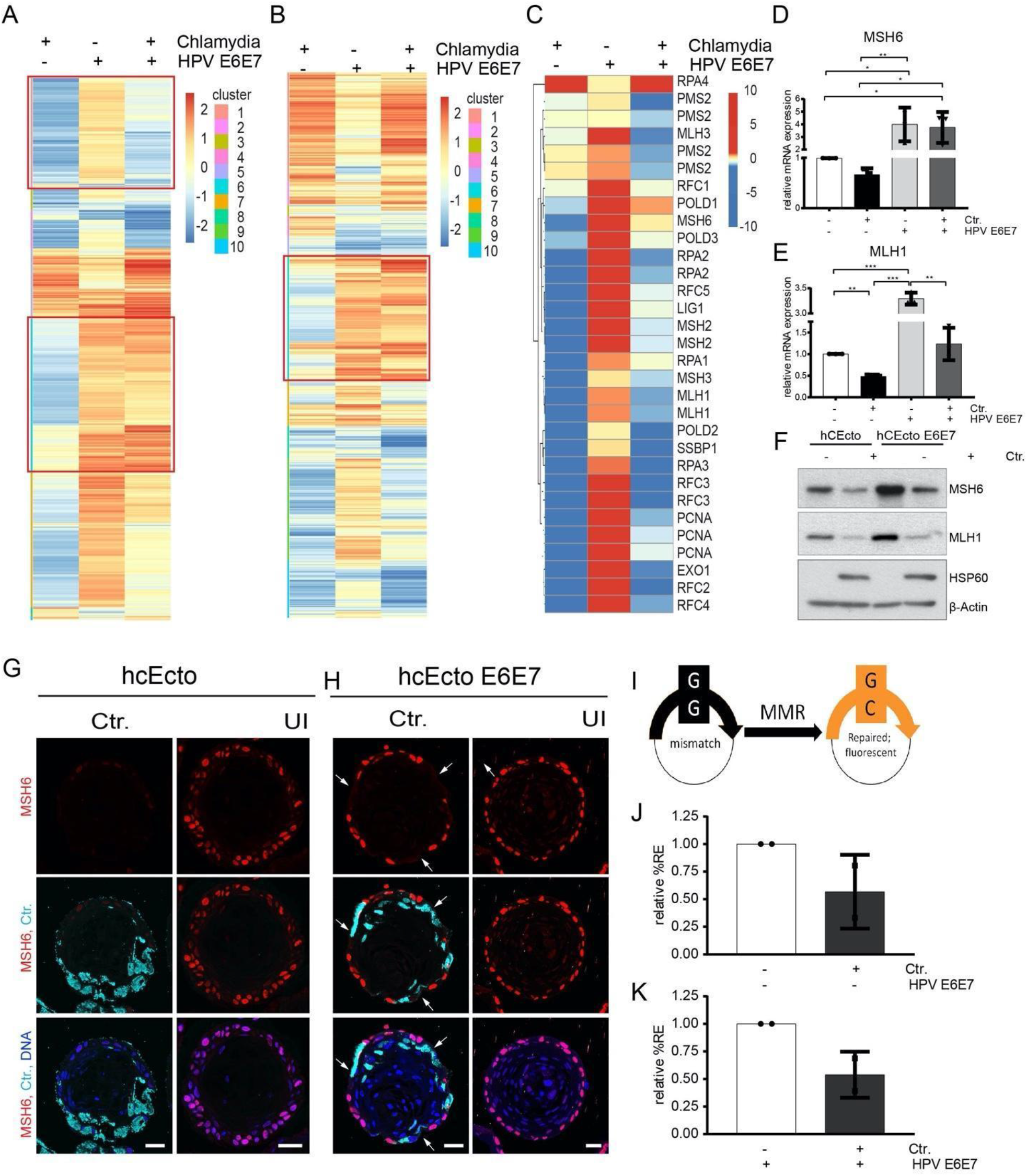
Inverse regulation of mismatch repair pathway by *C. trachomatis* and HPV E6E7 in human ectocervical epithelial stem cells and organoids. (A) Heatmap showing all DREAM target genes (Fischer, Grossmann et al. 2016) arranged into ten clusters based on their expression pattern between comparisons among the differentially regulated genes in hCEcto and hCEcto E6E7 2D stem cells with or without Ctr infection for 48 h. Red frames highlight the cluster containing mismatch repair (MMR) genes. The color bar depicts Z-scored expression values. (B) Heatmap showing all E2F target genes arranged into ten clusters based on their expression pattern between comparisons among the differentially regulated genes in hCEcto and hCEcto E6E7 2D cells with or without Ctr infection for 48h. Red frames highlight the cluster containing MMR genes. The color bar depicts log2 FC. (C) Heatmap depicting the expression of MMR pathway genes demonstrating the opposite effect of Ctr and HPV16 E6E7 on their regulation. (D-E) hCEcto and hCEcto E6E7 2D stem cells were infected with Ctr for 48 h. Shown is the relative mRNA expression of MSH6 (D) and MLH1 (E) analyzed by qRT-PCR. Data represent means ±SDs from three biological replicates normalized to uninfected controls. **p<0.01, *p<0.05, Student’s t-test. (F) hCEcto and hCEcto E6E7 2D stem cells were infected with *Ctr* for 48h, and cell lysates were subjected to immunoblot analysis for indicated proteins and chlamydia HSP60 and β-actin as a loading control. Data represent four biological replicates. (G-H) Shown are the hCEcto (G) and hCEcto E6E7 (H) organoids with or without Ctr infection for 5 d and immunolabeled for Msh6, Ctr-MOMP, and DNA in blue. The arrow highlights the loss of Msh6 expression in hCEcto E6E7. Scale bar: 50 μm. (I-K) A reporter plasmid containing a G:G mismatch was used for fluorescence-based multiplex flowcytometric host cell reactivation assay (FM-HCR) to analyze MMR efficiency by the restoration of reporter protein expression. (I) Comparison of relative reporter expression [%RE] between uninfected and Ctr infected hCEcto 2D stem cells (I) and hCEcto E6E7 (J) cells at 24 hpi showing reduced mismatch repair efficiency after infection. Data represent means ±SDs from two independent experiments normalized to uninfected controls. (*p<0.05, **p<0.01, ***p<0.001). Student’s t-test.

Mutational signatures associated with specific mutational processes have been found in a variety of human cancers; two signatures attributed to defective DNA mismatch repair (Hause, Pritchard et al. 2016, Bonneville, Krook et al. 2017) are found in cervical cancer (Antill, Dowty et al. 2015, Le, Durham et al. 2017). Previously, we have shown that *C. trachomatis* induces oxidative damage, particularly 8-oxo-dG (Chumduri, Gurumurthy et al. 2013). However, MMR, which plays a crucial role in preventing mutations because of oxidative damage (Bridge, Rashid et al. 2014), is upregulated by HPV E6E7 but downregulated by *C. trachomatis* infection. *C. trachomatis* suppressed the HPV E6E7 induced MMR pathway (Figure 4C Figure 3K, Supplementary Figure 2G). We found most of the genes involved in MMR, including *MLH1*, *MSH2*, and *MSH6* that play a central role in mismatch detection and initiation of the MMR signaling process (Kunkel and Erie 2005), are strongly downregulated by *C. trachomatis* infection in both hCEcto stem cells and organoids (Supplementary Figure 3A). *C. trachomatis* infection of HPV E6E7 expressing cells leads to a suppression of HPV-induced MMR genes (Figure 4D-E). Corroborating the global transcriptomic data, MSH6 and MLH1 protein levels were upregulated by HPV E6E7 but reduced by Chlamydia in single and HPV coinfections (Figure 4F, Supplementary Figure S3B-C). We confirm that MMR protein downregulation in Chlamydia infection is not an artifact resulting from chlamydial protease-like activity factor (CPAF)-mediated post cell lysis degradation of these proteins as the control protein golgin-84 remains uncleaved. (Supplementary Figure S3D). Thus, Chlamydia suppresses MMR at both transcriptional and post-translational levels, and it can further suppress HPV-activated MMR.

Since organoids are complex epithelial structures composed of different cell populations and differentiation stages, we investigated the spatial distribution of the MMR protein MSH6 in normal and *C. trachomatis* infections. Confocal images of the organoid sections from uninfected or 5dpi revealed high expression of MSH6 in the nucleus of basal stem cells with higher intensities in hCEcto E6E7 organoids corroborating with the increased protein levels. Further, a complete loss of MSH6 was observed in the *C.trachomatis* infected hCEcto organoids (Figure 4G). Interestingly, in hCEcto E6E7 organoids, *C. trachomatis* infected cells and neighboring cells show reduced or complete loss of MSH6 signals (arrows) (Figure 4H). Next, we sought to analyze if the impaired MMR pathway culminates in altered MMR capacity within the *C. trachomatis* infected host cells. A functional assay employing a G:G mismatch containing reporter plasmid that expresses a nonfluorescent protein (mOrange) was used to investigate this (Nagel, Margulies et al. 2014). Efficient repair of this in vitro generated G:G mismatch restores the wild type cytosine in the transcribed strand of the plasmid leading to mOrange fluorescent protein expression (Figure 4I). To determine transfection efficiency, hCEcto and hCEcto E6E7 cells were simultaneously transfected with a G:G mismatch-containing reporter plasmid and an undamaged reporter plasmid (expressing mPlum) prior to *C. trachomatis* infection. Fluorescence intensity was measured 24hpi by flow cytometry in living cells. The data showed that *C. trachomatis* infection leads to impaired MMR repair efficiency in single and HPV E6E7 (co-) infections, reflected by less reporter expression (%RE) in infected samples (Figure 4J-K).

### E2F-p53 and proteasomal degradation diverges the transcriptional and post-translational regulation of MMR during coinfections

Further, we sought to gain a mechanistic understanding of how Chlamydia and HPV regulate the MMR pathway. MMR genes are regulated by E2F transcription factors (Polager, Kalma et al. 2002, Bracken, Ciro et al. 2004). There is extensive crosstalk between the Rb–E2F and *MDM2*–p53 pathways, and remarkably, both are defective in most human tumors, which underscores the crucial role of these pathways in regulating vital cellular decisions. Since *C. trachomatis* induces MDM2-mediated and proteasome-dependent p53 degradation (Gonzalez, Rother et al. 2014, Siegl, Prusty et al. 2014), we checked if proteasomal degradation might be the underlying mechanism involved in MMR suppression. Using well-established proteasome inhibitors MG-132 (Maki, Huibregtse et al. 1996) and lactacystin to inhibit cellular and Chlamydia protease CPAF (Zhong, Fan et al. 2001) we tested the expression levels of MMR transcripts. Next, we checked the effect of p53 stabilization on regulating MMR gene expression and protein levels using Nutlin-3a that inhibits the interaction between MDM2 and p53 to stabilize p53 (Vassilev, Vu et al. 2004). RT-qPCR analysis showed that Nutlin-3a and MG-132 treatment restored MMR gene expression after *C. trachomatis* infection (Figure 5A-B). Lactacystin treatment increased MLH1 expression moderately compared to the control, but not MSH6. However, treatment with all three inhibitors created an unfavorable environment for *C. trachomatis* growth within the host cells, shown by a significant reduction in inclusion size compared to untreated control (Supp. Figure 4A). Further, immunoblot analysis revealed that *C. trachomatis* induced p53 degradation is inhibited by Nutlin-3a and MG-132 treatment, while E2F1 degradation was reduced in all treatment conditions. Of note, the MMR proteins MLH1 and MSH6 were restored only by inhibiting proteasomal activity with MG-132 (Figure 5C). The data show that Chlamydia engages p53-independent proteasomal degradation to reduce E2F1 and MMR proteins, while restoration of E2F1 and p53 proteins rescues transcriptional suppression of MMR genes (Figure 5G).

**Figure 5:**
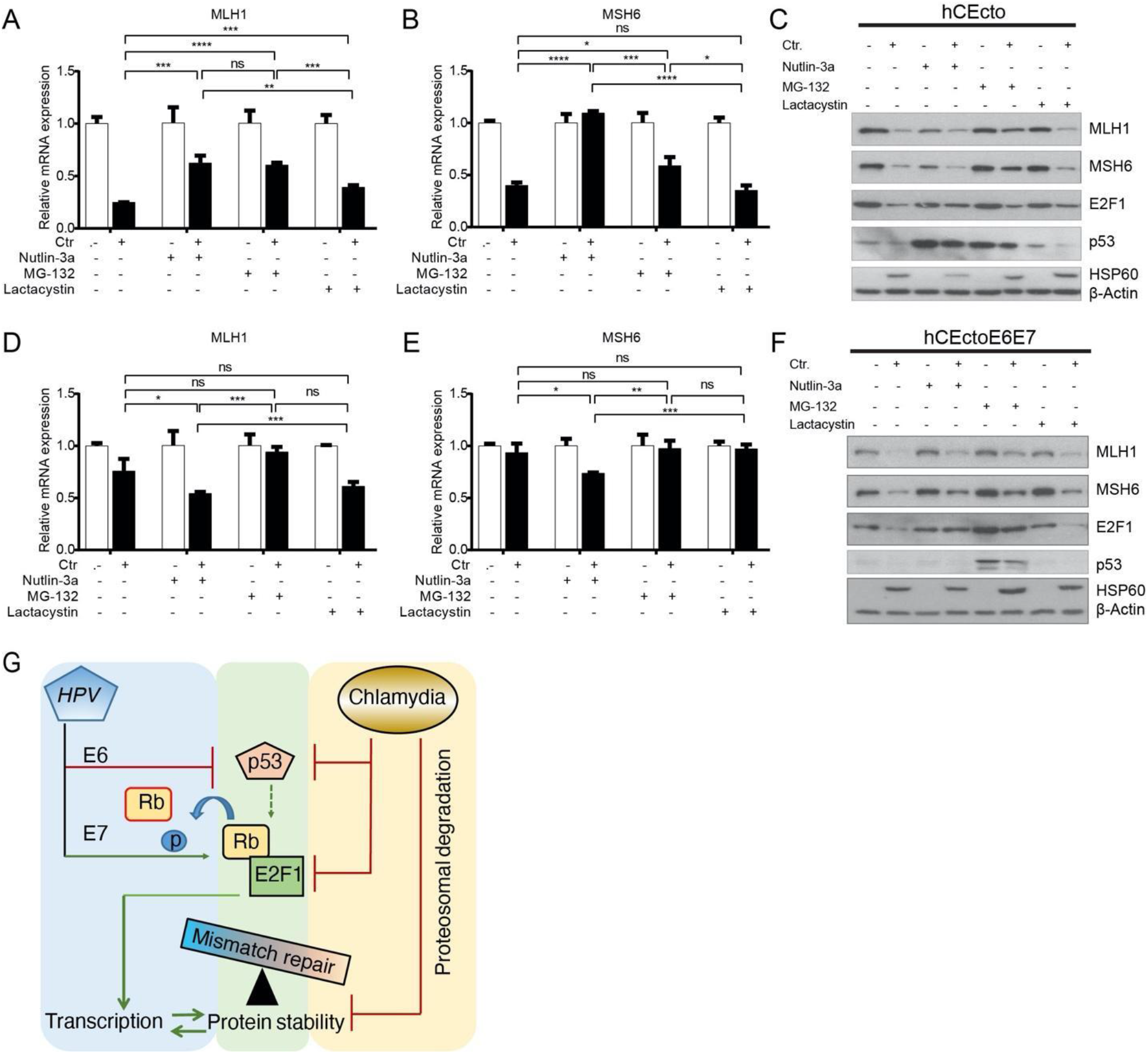
Chlamydia and HPV E6E7 coinfection regulates MMR distinctly at transcriptional and translational levels. hCEcto and hCEcto E6E7 2D stem cells were infected with Ctr for 48h with or without additional treatment with Nutlin-3a (10μM) or Lactacystine (10μM) from 2hpi or 24hpi with MG-132 (5μM). (A-B) Shown is the relative mRNA expression of MLH1(A) and MSH6 (B) analyzed by qRT-PCR from hCEcto cells. Data represent means ± SD of three replicates normalized to uninfected controls from a representative of two independent experiments. *p<0.05, **p<0.01, ***p<0.001, ****p<0.0001, ns, not significant. Student’s t-test. (C) Immunoblot analysis of lysates from hCEcto cells for indicated proteins and chlamydia HSP60 and β-actin as a loading control. (D-E) Shown is the relative mRNA expression of MLH1(D) and MSH6 (E) analyzed by qRT-PCR from hCEcto E6E7 cells. Data represent means ± SD of three replicates normalized to uninfected controls from a representative of two independent experiments. (F) Immunoblot analysis of lysates from hCEcto cells for indicated proteins and chlamydia HSP60 and β-actin as a loading control. Data from C and F represent two biological replicates. *p<0.05, **p<0.01, ***p<0.001, ns, not significant. Student’s t-test. (G) Model depicting the mechanistic regulation of MMR by HPV E6E7 and Chlamydia infections.

To further investigate how *C. trachomatis* regulates the MMR pathway in HPV E6E7 coinfections, hCEcto E6E7 stem cells were treated with Nutlin-3a, MG-132, and lactacystin with or without *C. trachomatis* infection for 48h. Like hCEcto cells, all treatment conditions reduced *C. trachomatis* infectivity as shown by significantly smaller inclusions than in control (Supplementary Figure 4B). In contrast to hCEcto cells, Nutlin-3a treatment of *C. trachomatis* infected hCEcto E6E7 cells shows further suppression of *MLH1* or *MSH6* gene expression (Figure 5D-E). However, proteasome inhibition by MG-132 led to a marginal rescue of MLH1 or MSH6 gene and protein expression. Further, Nutlin-3a and MG-132 treatment rescued the E2F1 protein expression. Notably, inhibition of proteasomal degradation by MG-132 stabilized p53, while Nutlin-3a and lactacystin treatment do not influence HPV-mediated degradation of p53 (Figure 5F). Together, proteasomal degradation is a crucial mechanism that distinctly regulates E2F-mediated MMR gene transcription and reduces p53, E2F, and MMR protein in coinfections (Figure 5G).

## Discussion

Insights into the coinfection of viruses and bacteria resulting in disease exacerbation are emerging (Griffiths, Pedersen et al. 2011, Kaul, Rathnasinghe et al. 2020). Mucosal surfaces of the female reproductive tract (FRT) play a central role in protecting the reproductive tract from infections while maintaining tissue homeostasis to prepare for successful fertilization and childbirth. Since it is critical to maintain sterile conditions in the upper FRT comprising the uterus, fallopian tube, and ovaries, the lower FRT, particularly the cervix, acts as a gatekeeper between the uterus and the vagina bearing the burden of defending against ascending infections. Despite this enormous importance of the cervical epithelium in maintaining women’s health and human reproduction, the lack of in vitro models that recapitulate the in vivo physiological epithelium has hampered the understanding of the single and coinfection processes that occur at the cervical epithelial barrier.

Our recently established human ectocervical organoid model that faithfully recapitulates the in vivo epithelial architecture of stratified squamous epithelium with its tissue-specific characteristics is a near-physiological in vitro model to investigate pathogenesis mechanisms (Chumduri, Gurumurthy et al. 2018, Chumduri, Gurumurthy et al. 2021). Since HPV is the etiological agent of cervical cancer development and often associated with simultaneous infections with *C. trachomatis* (Walboomers, Jacobs et al. 1999, Koskela, Anttila et al. 2000, Seraceni, Campisciano et al. 2016), the ectocervical organoids from HPV negative healthy donors are ideal tools to investigate the unexplored connection and the impact of HPV, *C. trachomatis* and coinfections on the host cells. The persistence of HPV triggered by integrating viral oncogenes E6 and E7 into host genome predisposes host cells for malignant transformation (Duensing, Lee et al. 2000, Filippova, Song et al. 2002, Yim and Park 2005). Our study simulated the impact of persistent HPV infection by integrating HPV16 E6E7 oncogenes into the human ectocervical epithelial stem cell genome. Notably, these human ectocervical stem cells containing HPV persistence developed the characteristics of CIN1. The organoids derived from these stem cells show increased epithelial regeneration by enhanced proliferation and differentiation, nuclear atypia with varied nuclear size, hyperkeratinization, and altered adherens junctions. However, they retain the capacity to self-renew, organize and differentiate to form mature squamous stratified organoids similar to healthy tissue with basal p63+ Ki67+ stem cells, p63+ Ki67-parabasal, and p63-Ki67-differentiated cells. In contrast, high grade lesions CIN2 and CIN3 tissues are characterized by the presence of Ki67+ and p63+ cells extending to all the stratified layers (Martens, Arends et al. 2004, Shirendeb, Hishikawa et al. 2009) indicating that integration of HPV E6E7 alone is not sufficient to break the stratified epithelial tissue patterning observed in high-grade lesions.

Despite Chlamydia being one of the four most frequent sexually transmitted pathogens, its infection process in complex ectocervical stratified epithelial tissue is unknown. This study systematically illustrated the temporal and spatial infection process in the multilayered ectocervical organoids recapitulating the physiological infections. Chlamydia can infect ectocervical stem cells and complete its biphasic life cycle. *C. trachomatis* infection propagates in stem cells, spread to differentiated layers, and ultimately released into the lumen. Our previous study, using 3D trans-well based air-liquid interface cultures of the human squamous stratified ectocervical epithelium, demonstrated the ability of *C. trachomatis* to infect differentiated luminal cells and its progression towards the basal stem cell compartment by disruption of epithelial integrity and induction of epithelial to mesenchymal transition phenotype that might cause tissue scarring (Zadora, Chumduri et al. 2019). This indicates that the type of cell Chlamydia infects will decide the infection and pathogenesis outcome. Our study provides the first insights into the influence of HPV persistence on *C. trachomatis* development. Interestingly, HPV E6E7 slows down the *C. trachomatis* developmental life cycle by inhibiting the redifferentiation of RBs into EBs and induces persistence.

Our global transcriptomic analysis revealed that HPV E6E7 and *C. trachomatis* elicit distinct host cell transcriptional programs. While tumor necrosis factor (TNF) mediated immune response is upregulated by both pathogens, *C. trachomatis* distinctively elicit IL-17 and NF-kappa B signaling inflammatory response. Chlamydia also markedly interferes with oxidative phosphorylation, RNA regulation, RNA processing, and upregulated MAPK pathways. Strikingly, superinfection of persistent HPV infected cells with *C. trachomatis* suppress a significant number of host genes induced by HPV E6E7, thus overwriting the host transcriptional program.

We found that HPV E6E7 activates the E2F family of transcription factors and their downstream-regulated genes involved in several cellular and genome surveillance pathways. Interestingly, Chlamydia superinfection of these cells suppresses E2F mediated regulation of DNA repair pathways and quiescence, despite inducing DNA damage through the production of reactive oxygen species (Chumduri, Gurumurthy et al. 2013). Multiple signatures of mutational processes are identified in cervical cancer samples, with two signatures associated with defective MMR (https://cancer.sanger.ac.uk/cosmic/signatures_v2). In line with previous observations (Moody and Laimins 2009, Spriggs and Laimins 2017), HPV is found to increase the MMR repair pathway. In contrast, *C. trachomatis* inhibits MMR at both the transcriptional and post-translational levels and further, suppresses the HPV-induced MMR activities. The *C. trachomatis* infected cells have a reduced ability to repair mismatches. Our data revealed that *C. trachomatis* inhibits MMR gene expression by proteasomal degradation of transcriptional factor E2F1. Suppressed MMR pathway coupled with the increased DNA damage induced by *C. trachomatis* could accelerate the mutagenic activity in the context of proliferation signals. The impact of HPV and Chlamydia’s co-persistence in a long-lived stem cell could, thus, be deleterious to cellular and genomic stability and promote neoplastic progression.

Our cervical 3D organoids provide the much-needed near-physiological in vitro models to investigate various facets of cervix biology, including the influence of FRT infections and coinfections and their molecular mechanisms. The long-term culturability of these organoids and the ability to genetically manipulate them for the first time opens avenues of approach to investigate the initiation, progression and outcome of chronic infections in an authentic preclinical setting. By utilizing this powerful development, we show here the complex tripartite interactions of epithelial tissue, viral and bacterial coinfection, and the impact on host cell response and fate. Thus, this study goes beyond state-of-the-art and shows how multiple sequential infections contribute to pathogenesis, and drive cells towards cancer progression.

## Acknowledgments

The authors would like to thank late Jörg Angermann and Christiane Dimmler for technical help, Ina Wagner for the microarrays, and Diane Schad for help with graphics. The work was mainly supported by the Max Planck Institute for Infection Biology and the University of Würzburg. C.C. acknowledges funding from DFG GRK2157. T.F.M acknowledges funding from BMBF via the Infect-ERA program CINOCA, Z.N. acknowledges funding from U01ES029520 and P30ES00002.

## Author Contributions

CC designed and led the study. T.F.M. provided the infrastructure and guidance. S.K., R.K.G., M.D., C.C., performed experimental work and analyzed the data. G.G., V.B. obtained electron micrographs, H.B., H.-J.M., S.K., R.K.G., and C.C., performed microarray analysis. M.M. provided human samples. Z.N. provided the MMR reporter plasmids. S.K., R.K.G., and C.C. wrote the manuscript. All authors read the manuscript and provided feedback.

---

**Supplementary Figure S1:**
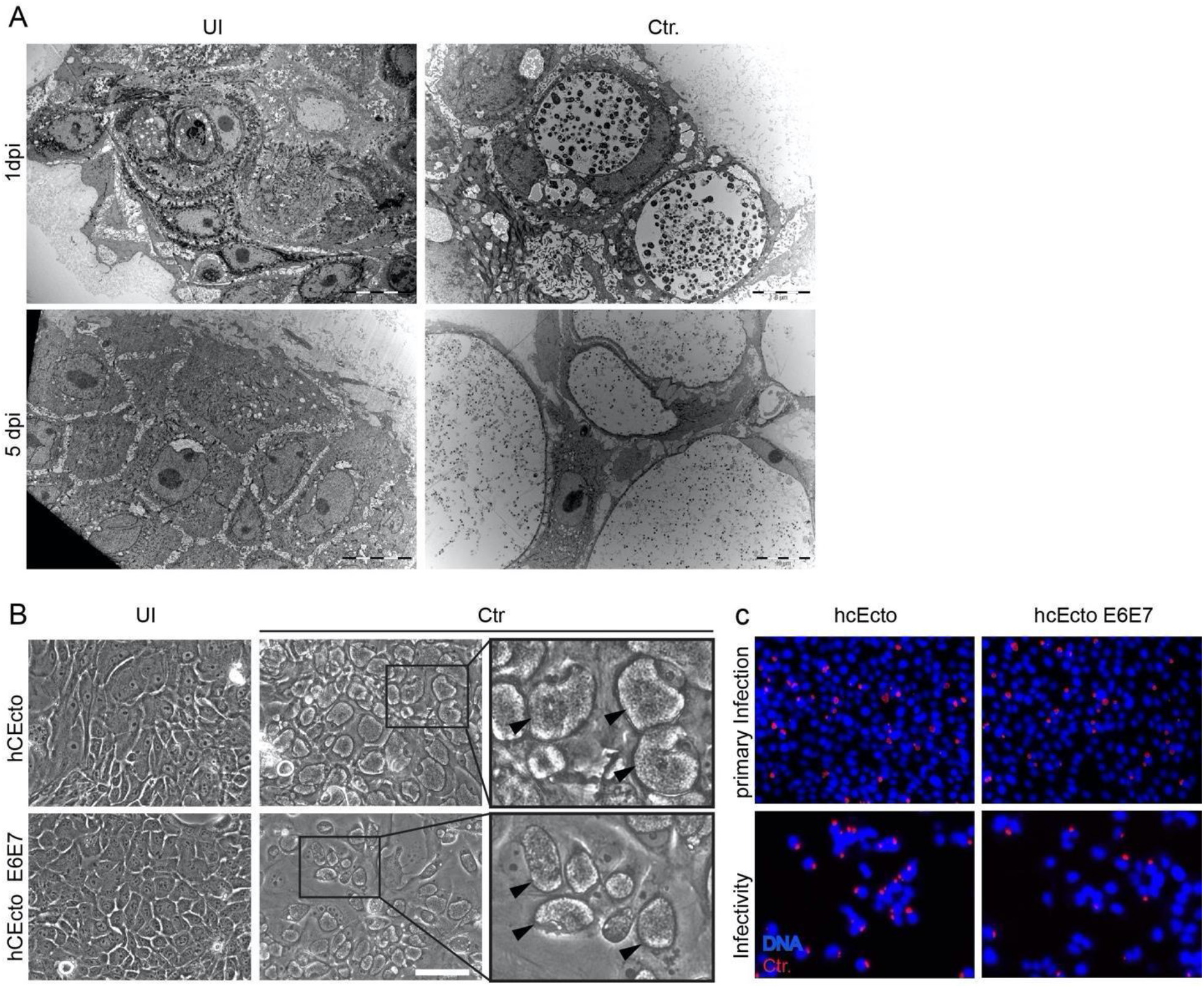
Ectocervical organoids as models for *C. trachomatis* and HPV coinfection studies. (A) Transmission electron micrographs of hCEcto organoids at 1 and 5dp Ctr infection. Scale bar: 10 μm (5dpi), 5 μm (1dpi). (B) Representative phase-contrast images of uninfected (UI) and Ctr-infected hCEcto and hCEcto E6E7 2D stem cells at 48 hpi, arrowheads in inserts show inclusions. Scale bar 150 μm. (C) Representative fluorescent images of Ctr primary infection and infectivity of hCEcto and hCEcto E6E7 cells. Images were taken with an automated microscope at 10x magnification.

**Supplementary Figure S2:**
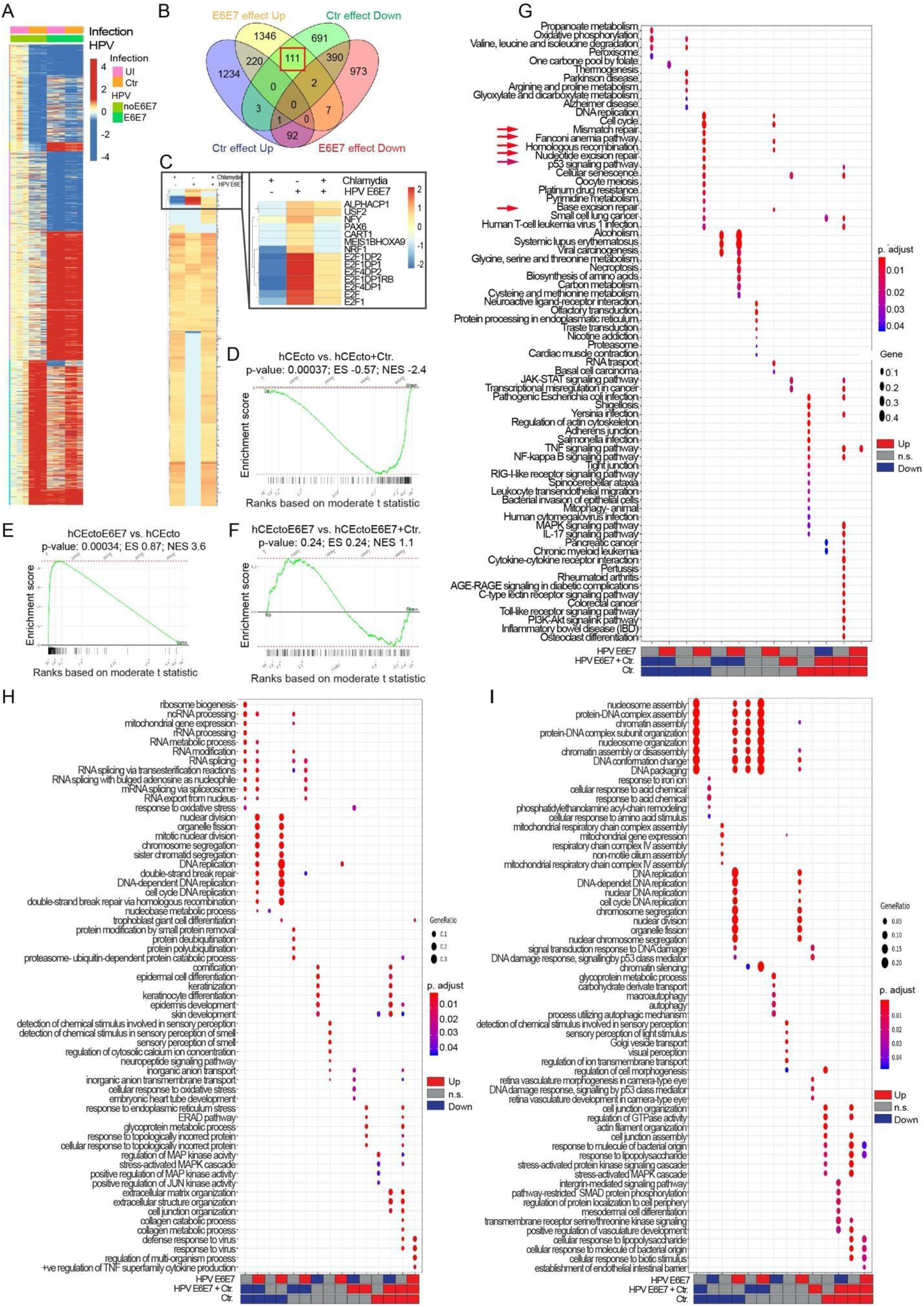
Host transcriptional response in human ectocervical organoids of *C. trachomatis* and HPV16 E6E7 and coinfection. (A) Heatmap of differentially regulated genes in hCEcto and hCEcto E6E7 organoids with or without Ctr infection for 5 d with p-value< 0.05; log2 FC >1; log2 FC < −1 between any of the conditions from three replicates. The color bar depicts expression values after subtracting the mean of UI/noE6E7 samples and dividing by SD of each probe. (B) Heatmap showing GSEA enrichment (-log10(p-value)) of genes that share Cis-regulatory motifs for transcription factors (stated in the right column). Significant enrichment of genes regulated by E2F1 transcription factor are upregulated by HPV E6E7 expression while being downregulated by single Ctr infection and in coinfection. (C) Venn diagram comparing genes significantly (log2 FC ≥ 2, p-value<0.05) up or down-regulated after Ctr infection or HPV E6E7 expression in hCEcto organoids. Highlighted in the red box are a subset of genes oppositely regulated by both pathogens, upregulated by HPV E6E7 and downregulated by Ctr. (D-F) GSEA analysis on differential gene expression comparisons of hCEcto vs hCEcto+Ctr (D), hCEcto vs hCEcto E6E7 (E), hCEcto E6E7 vs hCEcto E6E7+Ctr (F) organoids, identified E2F transcription factor target gene to be significantly downregulated after Ctr infection (D), upregulated upon HPV E6E7 expression (E) and suppression of HPV E6E7 induced effects upon coinfection with Ctr (F). (G) Dot plot showing significantly up or down-regulated KEGG pathways among the differentially expressed genes from hCEcto and hCEcto E6E7 organoids with or without Ctr infection categorized into distinct groups as indicated. Red arrows highlight DNA repair pathways. Dot diameter refers to the gene ratio within a group. Fill color depicts the p. adjust, Adjusted p-value. (H-I) Gene ontology (GO) terms associated with differentially expressed genes from hCEcto and hCEcto E6E7 organoids with or without Ctr infection categorized into distinct groups as indicated from 2D stem cells (H) and 3D organoids (I).

**Supplementary Figure S3:**
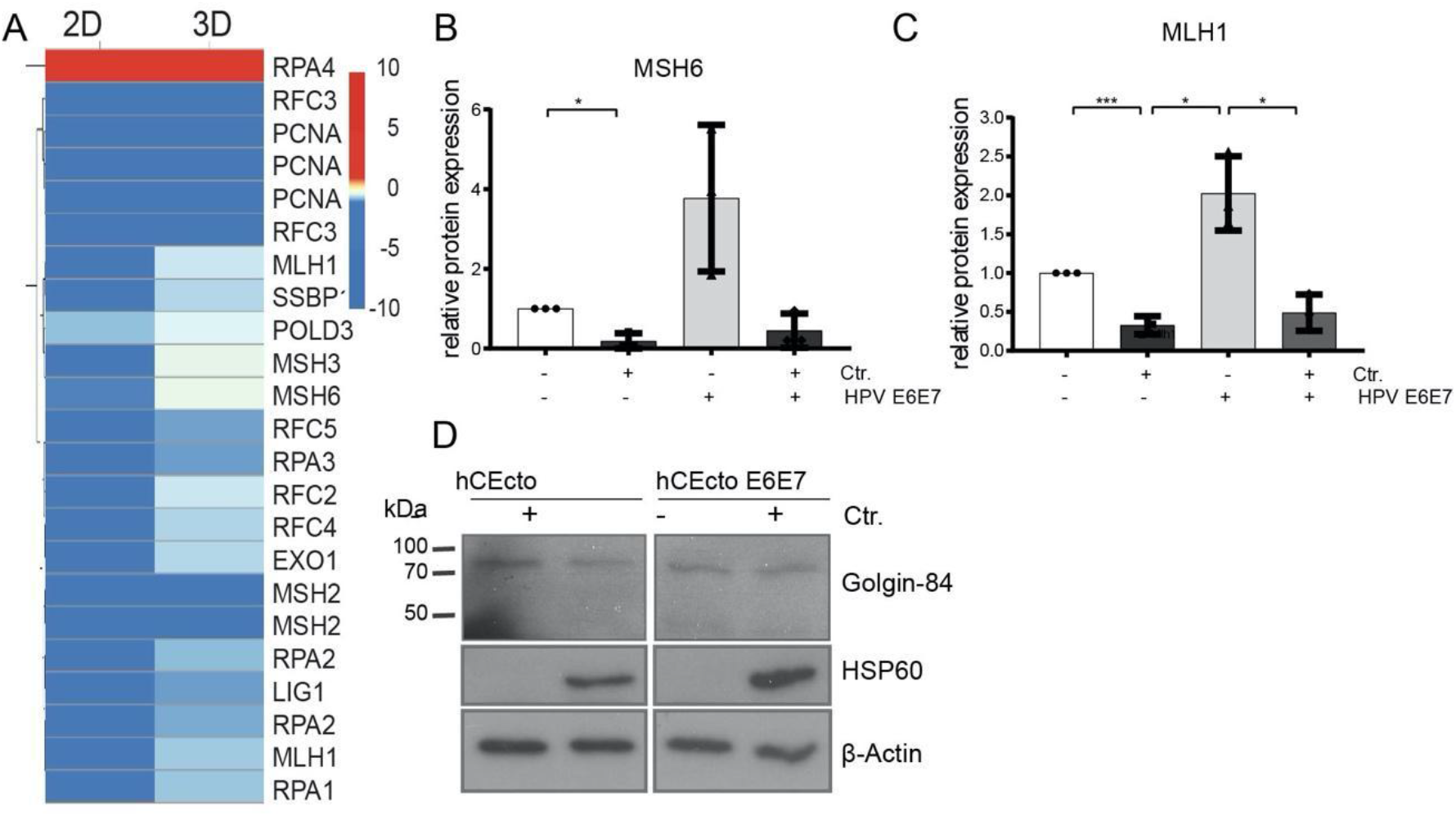
Chlamydia suppresses MMR in human ectocervical 2D stem cells and organoids. (A) Heatmap depicting the differential expression of MMR pathway genes from hCEcto 2D stem cells and organoids infected with Ctr for 48 h or 5 d, respectively. The color bar depicts log2 FC. (B-C) Shown are the densitometry quantification of MSH6 (B) and MLH1 (C) from immunoblots shown in Figure 4F. Densitometry values for MSH6 and MLH1 immunoblots were normalized to the β-actin values, and data representing the relative fold change compared to an uninfected and untreated control are shown. *p<0.05, **p<0.01, ***p<0.001), Student’s t-test. (D) hCEcto and hCEcto E6E7 2D stem cells were infected with Ctr for 48 h, and cell lysates were subjected to immunoblot analysis for indicated proteins and chlamydia HSP60 and β-actin as a loading control. kDa, kilo Dalton.

**Supplementary Figure S4:**
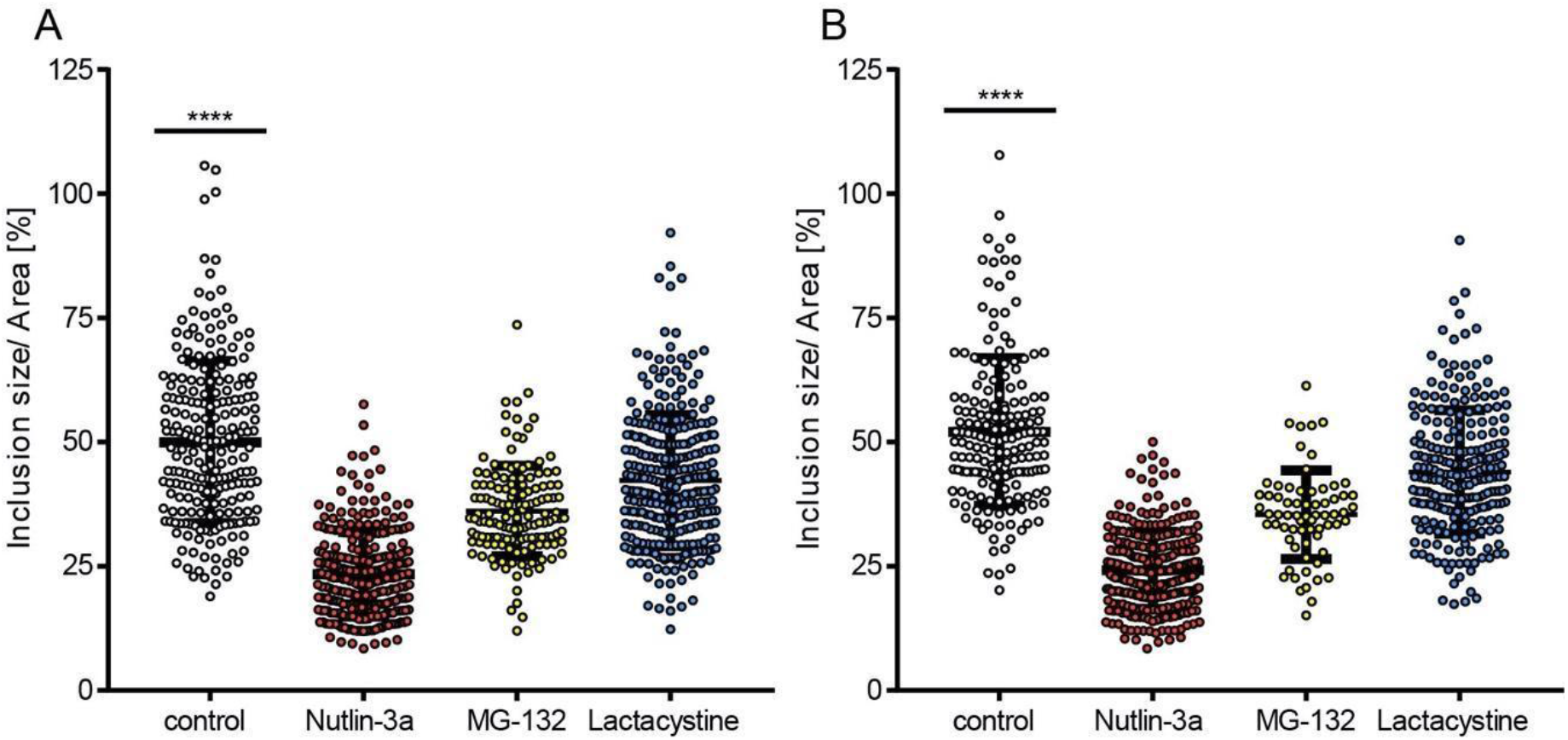
Impact of inhibition of proteasomal degradation and p53-MDM2 axis on *C. trachomatis* development. hCEcto and hCEcto E6E7 2D stem cells were infected with Ctr for 48h with or without additional treatment with Nutlin-3a (10μM) or Lactacystine (10μM) from 2 hpi or 24 hpi with MG-132 (5μM). Quantification of inclusion size by ImageJ from hCEcto (n ≥ 66 inclusions) (A) and hCEcto E6E7 (n ≥ 144 inclusions) (B). Data represent means ± SD. ****p<0.0001, Student’s t-test.

## Material and Methods

### Patient material

Human cervical samples were provided by the Department of Gynecology, Charité University Hospital, and August-Viktoria Klinikum, Berlin. Usage for scientific research was approved by their ethics committee (EA1/059/15); informed consent was obtained from all subjects. The study complies with all relevant ethical regulations regarding research involving human participants. Biopsies were sourced from standard surgical procedures. Only anatomically normal tissue biopsies from anonymous donors were processed within 2-3 h after removal.

### Human ectocervical primary cell isolation, cultivation, and maintenance

Isolation of epithelial stem cells from the human cervix, cultivation, and maintenance in 2D and organoids was performed as described in (Chumduri, Gurumurthy et al. 2018, Chumduri, Gurumurthy et al. 2021). In brief, tissue biopsies were washed, and the epithelial progenitors were isolated by mincing and enzymatic digestion using collagenase type II (0.5mg/mL) (Calbiochem, # 234155) (2.5 h, 37 °C). The dissociated cells were pelleted (5 min, 1,000g, 4 °C), resuspended in TrypLE (Gibco, # 12604021), and incubated in a shaker (15 mins, 37°). Primary cells were expanded in collagen-coated (Sigma, # C3867) tissue culture flasks until they reached 70-80% confluence in human ectocervical primary cell medium (Advanced DMEM/F-12 (Invitrogen, 12634) supplemented with 12 mM HEPES, 1% GlutaMax, 1% B27, 1% N2, 10 ng/mL human EGF (Invitrogen, 15630-056, 35050-038, 17504-044, 17502048, PHG0311), 0.5 μg/mL hydrocortisone (Sigma, H0888-1G), 100 ng/mL human noggin (Peprotech, 120-10C), 100 ng/mL human FGF-10 (Peprotech, 100-26-25), 1.25 mM N-acetyl-L-cysteine (Sigma, A9165-5G), 2 μM TGF-β receptor kinase Inhibitor IV (SB431542, Calbiochem), 10 μM ROCK inhibitor (Y-27632) (Sigma, Y0503), 10 mM nicotinamide (Sigma, N0636), 10 μM forskolin (Sigma, F6886)) before seeding ~20,000 cells/50 μL in Matrigel (BD, # 356231) for culturing organoids or for maintenance of 2D stem cells on lethally irradiated J2-3T3 fibroblast feeder cells. For propagation, 2D stem cells were reseeded onto freshly irradiated J2-3T3 every 1 to 2 weeks at a 1:5 ratio. Ectocervical organoids were split every 2 weeks at a ratio of 1:5 using enzymatically splitting with TrypLE and mechanical fragmentation with a fire-polished glass Pasteur pipette by vigorous pipetting.

For 2D infection experiments of human ectocervical epithelial (hCEcto) stem cells, the cells were subjected to differential trypsinization to separate fibroblasts from epithelial cells, and only epithelial cells were seeded on collagen-coated cell culture plates. Plates were coated with collagen type I at diluted 1:100 ratio with PBS (solution from rat tail) (Sigma-Aldrich, # C3867) for 1 h at 37°C. Cells were incubated at 37°C 24h before infection.

### Cell lines

HeLa cells (ATCC® CCL-2™), End1 E6E7 cells (ATCC® CRL-2615™), 293T cells (Invitrogen-10938-025), and J2-3T3 cells (mouse embryo) (kind gift from Craig Meyers) were maintained in HEPES-buffered Dulbecco’s modified Eagle’s medium (DMEM) (Gibco, 10938-025) supplemented with 10% fetal calf serum (FCS) (Biochrome, S0115), 2 mM glutamine (Gibco, 25030081), and 1mM sodium pyruvate (Sigma, S8636), at 37°C in a humidified incubator containing 5% CO_2_.

### *C. trachomatis* infections of human ectocervical stem cell

*C. trachomatis* L2 (ATCC, # VR-902B) stocks were prepared as described earlier (Gurumurthy, Maurer et al. 2010, Chumduri 2018). *C. trachomatis* infection experiments in 2D stem cells were performed at a multiplicity of infection (MOI) of 5 in ectocervical primary cell medium. The medium was exchanged at 2 h post-infection (hpi) and cells were grown at 35°C in 5% CO_2_ in a humidified incubator for 24 h up to 48 h post-infection (hpi) and treated with chemical compounds depending on the experiment.

### *C. trachomatis* infections of human ectocervical organoids

For 3D ectocervical organoid *C. trachomatis* infection experiments, human ectocervical organoids were grown for five days in a 24-well plate at 37°C in 5% CO_2_ in human ectocervical primary cell medium without antibiotics. Then, media was removed, and Matrigel was dissolved by adding 1mL ice-cold ADF and pipetting up and down. The resuspended organoids were divided into two 15 mL tubes for infection and mock control. To release organoids completely from Matrigel, another 4mL of ice-cold ADF was added to each tube. After centrifugation at 300g for 5 min at 4 °C, the supernatant was discarded, resuspend the pellet in 250 μL medium containing diluted Ctr stock, to yield the approximate MOI of 10. Organoids were incubated for 2 h at 35°C with 5% CO_2_ with gentle shaking at 100 rpm. After infection, organoids were centrifuged at 300g for 5 min, and the pellet was resuspended in Matrigel (50 μL/ well) and kept at 35°C with 5% CO_2_ in a humidified incubator. Once Matrigel was polymerized, a fresh human ectocervical primary cell medium was added, and the infection was allowed to proceed until the desired time.

### Infectivity assay

hCEcto stem cells grown in collagen-coated six-well plates were infected with *Ctr-* L2 (MOI=1) for 48h and then scraped and transferred into a 15 mL tube containing sterile glass beads and lysed by vortexing for 4 min at full speed. To determine the infectivity from organoid infection experiments, Matrigel was first dissolved with ice-cold ADF, and the suspension was centrifuged (5 min at 300g, 4 °C). Then the supernatant was discarded, and the pelleted organoids were resuspended in 1 mL of DMEM supplemented with 5% FCS (infectivity medium) and 1 mL of sterile glass beads followed by 4 min vortexing at full speed to lyse the cells. The lysates’ dilutions from 2D stem cells and organoids were then processed similarly by transferring onto HeLa cells seeded 1 day before in 96-well plates and incubated for 2 h. The lysates were discarded, an infectivity medium was added, and cells were incubated overnight at 35°C with 5% CO_2_. Cells were fixed with ice-cold methanol 200 μL/well overnight at 4 °C and immunostained with *Ctr*-major outer membrane protein (*Ctr*-MOMP KK12) specific antibody and Cy3 labeled secondary antibody. Host cell nuclei were stained with Hoechst. The number of chlamydial inclusions, inclusion size, and the number of host cells were analyzed with an automated microscope and ScanR Analysis Software (Olympus Soft Imaging Solutions) as previously described (Gurumurthy, Maurer et al. 2010).

### Antibodies and Chemicals

Antibodies and chemicals were obtained from the following sources: mouse-anti-p63 (4A4) (1:200, Abcam, # ab735), mouse-anti-E-Cadherin (1:100, BD Biosciences, # 610181), rabbit-anti-Ki67 (SP6) (1:200, Abcam, # ab16667), rabbit-anti-Cytokeratin 5-Alexa488 (1:300, Abcam, # ab193894), rabbit-anti-Cytokeratin 8 (1:200, Abcam, # ab59400), rabbit-anti-Loricrin (1:50, Abcam, # ab85679), mouse-anti-phospho γH2AX (Ser139) (1:500, Millipore, #05636), rabbit-anti-Msh6 (EPR3945) (1:1000, Abcam, # ab92471), mouse-anti-Msh2 (3A2B8C) (1:1000, Abcam, # ab52266), rabbit-anti-Mlh1 (EPR3894) (1:10000, Abcam, # ab92312), mouse-anti-β-Actin (1:10000, Sigma, # 014M4759), mouse-anti-E2F1 (8G9) (1:250, Abcam, # ab 135251), mouse-anti-Rb (4H1) (1:2000, Cell Signaling, # 9309), rabbit-anti-pRb (Ser807/811) (1:1000, Cell Signaling, # 9308), mouse-anti-p53 (DO-1) (1:500, Santa Cruz, # sc-126), mouse-anti-Chlamydia Hsp60 (A57-E4) (1:500, Enzo Life Sciences, # ALX-804-071-R100 and 1:1000 GeneTex, # GTX25486) and *goat-anti-Chlamydia* major outer membrane protein (MOMP) (1:500, AB Biotec, # 804), Mouse monoclonal species-specific KK-12 IgG2a *Ctr* (anti-MOMP) (1:10000, D. Grayston, University of Washington, Seattle, WA, USA). Secondary antibodies used for immunofluorescence were donkey-anti-goat Alexa Fluor 488 (Jackson ImmunoResearch, # 705-545-147), donkey-anti-rabbit-Cy3 (Jackson ImmunoResearch, # 711-166-152), donkey-anti-mouse Alexa Fluor 647 (Jackson ImmunoResearch, # 715-605-150), goat anti-mouse-Cy3 (Dianova, # 115-165-006), donkey-a-goat-Dylight 647 (Dianova, # 705-605-003) (1:150) and donkey-anti-mouse Alexa Fluor 488 (Dianova, # 715-454-151). Hoechst (1:2000, Sigma, # B2261) and Draq5 (1:1000, Thermo Scientific, # 62252) were used to label DNA. Secondary antibodies conjugated to horseradish peroxidase (HRP) were purchased from Amersham Biosciencesa and used at 1:2000 dilution. Following chemicals were obtained from Sigma-Aldrich: Proteasome inhibitor MG132 (# M7449), Proteasome and chlamydial protease-like activity factor (CPAF) inhibitor Lactacystin (# L6785), and tumor suppressor p53 (TP53) stabilizer and MDM2 inhibitor Nutlin-3a (# N6287).

### RNA and DNA isolation

RNA and DNA were isolated using the Allprep and RNeasy Mini Kit (Qiagen, # 80204, # 74104) according to the manufacturer’s protocol.

### Polymerase chain reaction (PCR) and agarose gel electrophoresis

The presence of genes encoding for human HPV16 E6E7 and HPV18 E6E7 was detected using the PCR. The PCR mixture contained 100 ng DNA template, 0.5 μL forward primer (10 μM), 0.5 μl reverse primer (10 μM), 0.5 μl dNTPs (10 mM) (Thermo Scientific, # R0182), 2.5 μL 10x NEB buffer (BioLabs, # B9004S), 1 μL NEB Taq polymerase (5000U/mL) (BioLabs, # M0267S), 1 μL MgCl_2_ (25 mM) (Promega, # A351H) and the reaction was made up to 25 μL with H_2_O. The PCR cycling conditions were as follows: initial denature of 5 min at 95°C followed by 35 cycles at 95°C for 30s, at 55°C for 30s and at 72°C for 1 min. The final extension was at 72°C for 5 min. PCR products were directly processed using agarose gel electrophoresis. DNA fragments were separated in 1.5% agarose (Biozym, #840004) in 0.5% v/v TBE buffer supplemented with ethidium bromide solution at 120V for 60 min. Up to 15 μL of PCR product was loaded into the agarose gel using 6x DNA loading buffer (Fermentas, # R0611). Following primers were used: GAPDH-Forward: 5’-GGTATCGTGGAAGGACTCATGAC-3’, Reverse: 5’-ATGCCAGTGAGCTTCCCGTTCAG-3’; HPV16-Forward: 5’-AGCTGTCATTTAATTGCTCATAACAGTA-3’, Reverse: 5’-TGTGTCCTGAAGAAAAGCAAAGAC-3’; HVP18-Forward: 5’-CGAACCACAACGTCACACAAT-3’, Reverse: 5’-GCTTACTGCTGGGATGCACA-3’. The primers were purchased from Sigma-Aldrich and diluted to 10 μM.

### Quantitative real-time polymerase chain reaction (qRT-PCR) Analysis

Relative RNA levels were determined by RT-qPCR using Power SYBR® Green RNA-to-CT™ 1-Step Kit (Thermo Fisher, # 4389986), StepOnePlus ™ Real-Time PCR System (Applied Biosystems TM), and StepOneTM Software (v2.3, Applied Biosystems). The reaction mixture contained 25ng RNA, 12.5 μl SYBR® green mix, 1.84 μl H2O, 0.16 μl Reverse Transcriptase enzyme mix and 0.5 μl Primer mix (10 μM) and was subjected to the following PCR cycler program: 30 min at 48° C; 10 min at 95° C; followed by 40 cycles of 15s at 95° C and 60s at 60° C. The relative expression levels of all genes were determined by normalizing to mRNA expression of housekeeping gene Valosin Containing Protein (VCP). All samples were measured as triplicates. Following primers were used: MSH6-Forward: 5’-CCCCACCAGTTGTGACTTCT-3’, Reverse: 5’-TGTTGGGCTGTCATCAAAAA-3’; MSH2-Forward: 5’-TCTTCGTTCTGACTTCTCCAAG-3’, Reverse: 5’-ATCTGTTTGCCAGGGTCCA-3’; MLH1-Forward: 5’-TGAGCAGGGACATGAGGTTCTC-3’, Reverse: 5’-ACTAAGCTTGGTGGTGTTGAG-3’; VCP-Forward: 5’-AGCATTGACCCAGCTCTACG-3’, Reverse: 5’-TCTCATTGGCTACCTGTTCCAG-3’; E2F1-Forward: 5’-AGATGGTTATGGTGATCAAAGCC-3’, Reverse: 5’-ATCTGAAAGTTCTCCGAAGAGTCC-3’; RB-Forward: 5’-CTCTCGTCAGGCTTGAGTTTG-3’, Reverse: 5’-GACATCTCATCTAGGTCAACTGC-3’; TP53-Forward: 5’-CCTCCTCAGCATCTTATCCGA-3’, Reverse: 5’-TGGTACAGTCAGAGCCAACCTC-3’. The primers were purchased from Sigma-Aldrich and diluted to 10 μM. The melting temperature was determined to be 60°C for all primers.

### SDS-PAGE and Western blotting

Total cellular extracts from cells grown in six-well plates treated as per experimental requirements were prepared by incubating cells in 200-300 μL of SDS lysis buffer (3% 2-mercaptoethanol, 20% glycerin, 0.05% bromophenol blue (AppliChem, # A2331,0005), 3% SDS). Cell lysates were collected and boiled at 95°C for 10 min. Samples were stored at −20°C until required. SDS-PAGE and Western blotting were performed as described earlier (Gurumurthy, Maurer et al. 2010).

### Generation of HPV16 E6E7 expressing hCEcto primary stem cells by lentiviral manipulation

Replication-deficient lentiviral particles carrying HPV16 E6E7 or Luciferase gene were produced by transfection of 293T cells with pLXSN HPV16 E6E7, (Addgene, #52394) or pSRE-Luciferase plasmid from ATCC (# MBA-120) together with packaging plasmids pMD2.G (Addgene, #12259), psPAX, (Addgene, #212260) and Fugene6 (Promega, #E2691) diluted in Opti-MEM™ (Gibco, # 31985088). For a 10 cm dish, 15.6 μl Fugene6 was mixed with 192.4 μl OPTI-MEM and added to 2.6 μg lentiviral target plasmid, 1.95 μg psPAX2 packaging vector, and 0.65 μg pMD.2G (VSVG) envelope vector diluted in 52 μl OPTI-MEM. The DNA-Fugene Mix was incubated for 20-30 min at RT before it was dropwise added to the 293T cells. Cells were incubated with the mix for 12-15h at 37°C and 5% CO_2_ before the medium was replaced with fresh DMEM. Two days post-transfection lentiviral particles present in the medium were harvested, filtered (0.45 μm), and used for hCEcto cell transduction. For transduction, hCEcto stem cells were grown on collagen-coated six well-plate to 50% confluence before they were treated with virus solution and Polybrene (1μg/mL) (Sigma-Aldrich, # H9268) overnight at 37°C. When cells reached ~90% confluence, they were selected by 0.5 μg/mL Blasticidin (Gibco) treatment.

### Immunofluorescence histochemistry

Organoids and human tissue were fixed with 3.7% paraformaldehyde (PFA, Sigma-Aldrich, # 4441244), dehydrated, embedded in paraffin, sectioned, and stained for confocal imaging using Leica TCS SP8 (Leica Microsystems GmbH) as described previously (Chumduri 2018). hCEcto cells grown on collagen-coated coverslips in 2D were fixed with 3.7% paraformaldehyde for 30 min at RT. Cells were permeabilized and blocked with 2% Triton X100, 0.05% Tween 20, and 1% BSA (Biomol, # 01400.1) in PBS overnight at 4 °C. Primary antibodies were diluted in 1% BSA, 0.05% Tween 20 in PBS and incubated for 1 day at 4 °C followed by three washes in PSB-T (0.1% Tween 20 in PBS) and 1 h incubation with secondary antibodies diluted in 1% BSA, 0.05% Tween 20 in PBS along with Hoechst or Draq5. Before mounting with Mowiol, coverslips were washed three times with PBS-T and once with PBS. Images were acquired on a Leica TCS SP8 confocal microscope and were processed with ImageJ and Adobe Photoshop.

### Transmission electron microscopy

To analyze *C. trachomatis* development in human ectocervical epithelial cells, electron micrographs of infected and uninfected organoids and 2D stem cells were taken. Samples were prepared as described previously (Gurumurthy, Chumduri et al. 2014). Briefly, at different time points after *C. trachomatis* infection, cells were fixed with 2.5% glutaraldehyde for 2 days at 4 °C. Cells were postfixed with 0.5% osmium tetroxide and tannic acid, contrasted with uranyl acetate, dehydrated in ascending ethanol series, and infiltrated via styrene in epoxy resin. Samples were embedded in molds and allowed to polymerize at 70°C. Electron micrographs were taken from 60 nm sections, which were contrasted with lead citrate, with a Leo 906E transmission electron microscope operated at 100kV acceleration voltage (Zeiss, Oberkochen, DE) using a side-mounted digital camera (Morada, SIS-Olympus, Münster, DE).

### Mismatch DNA Repair Assay

#### Plasmids

Fluorescent and nonfluorescent reporter plasmids used for measuring DNA repair capacity of human primary epithelial cells under different conditions were kindly provided by Dr. Zachary Nagel (Harvard T.H. Chan School of Public Health).

#### Sample preparation

hCEcto 2D stem cells were propagated to achieve a confluence of ~70%. Cells were washed once with PBS, followed with TrypLE incubation for approximately 10-15 min at 37°C to obtain single cells.

#### Electroporation

For each sample, around 0.25 x 10^6^ cells were combined with a reporter plasmid mixture containing 0.5 ng of plasmid, that contains a G:G mismatch (G229C) (Table 1), in a total volume of 100μL/ reaction and electroporated using an Amaxa ^TM^ Basic Nucleofector ^TM^ Kit for Primary Mammalian Epithelial cells (Lonza, # VPI-1005) and the Nucleofector^TM^ 2b Device (Lonza) with the preset program T-20. Transfection efficiency in each experiment was measured by simultaneous transfection of fluorescent reporter plasmid mPlum. Following electroporation, 1 mL of human ectocervical primary cell medium was added, and the reaction mixture was divided into two tubes for infection and mock control. *C. trachomatis* infection was performed at a multiplicity of infection (MOI) of 5. The inoculum was added after Nucleofection to a tube, followed by 1 h incubation at 37°C with gentle shaking. The mock control was treated similarly. After infection, cells were pelleted by centrifugation (5 min at 1000 g; 4°C), resuspended in human ectocervical primary cell medium, plated onto collagen-coated plates, and incubated for 24h at 35 °C, 5% CO_2_ in a humidified incubator. Next, cells were washed with PBS, trypsinized with TrypLE for 15 min, and resuspended in 500 μL PBS containing Propidium iodide (PI, S Thermo Scientific Invitrogen^TM^, # P3566) stain and transferred to 75-mm Falcon tubes with Cell Strainer Caps (Fisher Scientific).

**Table 1:**
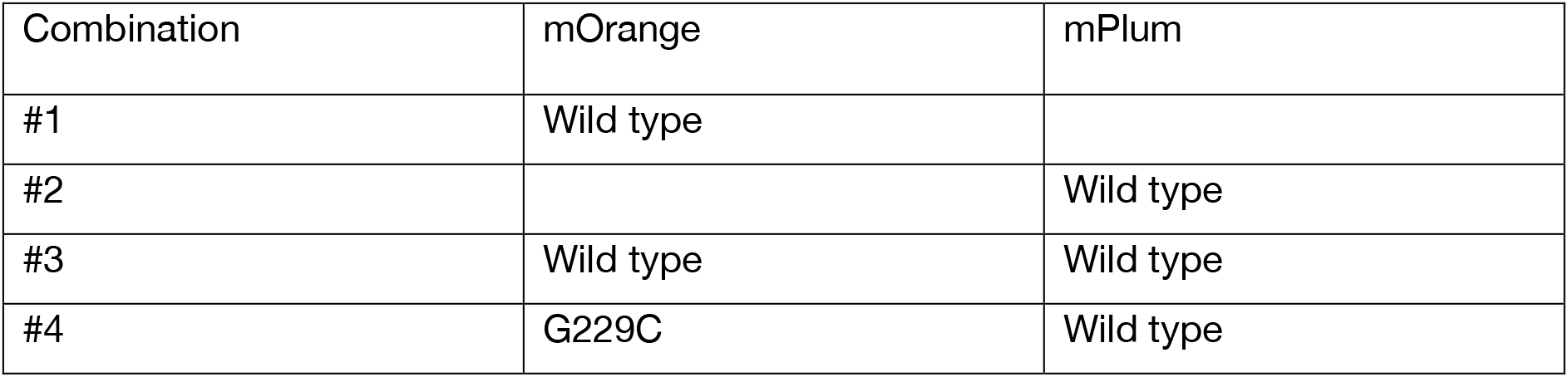
Combinations of reporter plasmids and types of DNA damage used in each experiment

| Combination | mOrange | mPlum |
| --- | --- | --- |
| #1 | Wild type |  |
| #2 |  | Wild type |
| #3 | Wild type | Wild type |
| #4 | G229C | Wild type |

#### Flow Cytometry

Cells were analyzed for fluorescence on a FACSymphony™ A5 (BDbiosciences) running the FACSDiva software (BDbiosciences). Any cell debris, doublets, or aggregates were excluded based on side-scatter and forward-scatter properties. The following fluorophores and their corresponding detectors were used: mOrange (505LP-580/20), mPlum (635LP-670/30), GFP (505LP-530/30). Compensation was set by using single-color controls. Data were quantified with FlowJo V10 (FlowJo, LLC). Equations used to calculate fluorescence reporter expression %R.E. have been previously described (Nagel, Margulies et al. 2014, Chaim, Gardner et al. 2017).

### Statistics

Results are presented as the mean ±SD (standard deviation). For Analysis, data sets were compared by unpaired or paired students t-test in GraphPad Prism 8 (GraphPad Software). Data are considered statistically significant with a p-value ≤ 0.05 unless otherwise specified.

### Microarray expression profiling and data analysis

2D stem cells and organoids were pelleted and resuspended in 1 mL Trizol® reagent (Invitrogen™, # 15596026). RNA was isolated according to the manufacturer’s protocol, and the quantity of RNA was measured using a NanoDrop 1000 UV-Vis spectrophotometer (Kisker), and quality was assessed by Agilent 2100 Bioanalyzer with an RNA Nano 6000 microfluidics kit (Agilent Technologies, # 5067-1511). All Microarray experiments were performed as single-color hybridizations. Total RNA was amplified and labeled with the Low Input Quick-Amp Labeling Kit (Agilent Technologies, # 5190-2305). In brief, mRNA was reverse transcribed and amplified using an oligo-dT-T7 promoter primer and labeled with cyanine 3-CTP. After precipitation, purification, and quantification, 0.75 μg labeled cRNA was fragmented and hybridized to custom whole-genome human 8 × 60K microarrays (Agilent-048908) according to the supplier’s protocol (Agilent Technologies). Scanning of microarrays was performed with 3 μm resolution (8×60K) using a G2565CA high-resolution laser microarray scanner (Agilent Technologies). Microarray image data were processed with the Image Analysis/Feature Extraction software G2567AA v. A.11.5.1.1 (Agilent Technologies) using default settings and the GE1_1105_Oct12 extraction protocol. The extracted single-color raw data files were background corrected, quantile normalized, and further analyzed for differential gene expression using Rstudio (Rstudio. Inc.). Microarray data from three independent infection experiments were combined. The signature of differentially expressed genes between *C. trachomatis* infected and uninfected hCEcto 2D stem cells or organoids with or without HPV16 E6E7 were selected from all genes in any of the four comparisons (hCEcto vs. hCEcto+*Ctr*; hCEcto vs. hCEcto+E6E7; hCEcto+E6E7 vs. hCEcto+E6E7+*Ctr*, hCEcto*+Ctr* vs. hCEcto+E6E7+*Ctr*). All genes differentially expressed with FDR < 5% (p-value ≤ 0.05) and log2 fold change <-1 or >1 for each comparison between any of those conditions were used.

For Venn diagram visualization with VENNY 2.1 [http://bioinfogp.cnb.csic.es/tools/venny/], statistically significant (p-value ≤ 0.05) up or down (FC > ±2) regulated genes after *C. trachomatis* infection or HPV16 E6E7 expression in primary ectocervical cells were chosen.

### Analysis of KEGG pathways and Gene ontology

For each of the conditions, the infected cells are compared to the non-infected baseline (i.e., condition with no *Ctr* and no E6E7) or E6E7 transduced cells. For both 2D stem cells and organoids, we classify each item (gene, gene set, or regulon) regarding its result in *C. trachomatis* alone, coinfection, or E6E7 alone in a single class (e.g., Down-Down-Up). Corresponding tables were generated, and for genes, an enrichment was run for each class. For further analysis, all subclasses with less than or equal to 20 genes and all that have conflicting (Up-Down) results for probes within the same gene were excluded.

### Gene Set Enrichment Analysis (GSEA)

A pre-ranked analysis using the R package fGSEA was used that should give similar results to pre-ranked analysis in standard GSEA. The t-statistics from comparisons of ectocervical 2D stem cells and organoids (hCEcto vs. hCEcto+*Ctr*; hCEcto vs. hCEcto+E6E7; hCEcto+E6E7 vs. hCEcto+E6E7+*Ctr* or hCEcto+*Ctr* vs. hCEcto+E6E7+*Ctr*) were used to rank probes and enrichment of MSigDB V6.1 gene sets [http://software.broadinstitute.org/gsea/msigdb] (h.all.v6.1.symbols.gmt; c2.all.v6.1.symbols.gmt; c3.all.v6.1.symbols.gmt; c5.bp.v6.1.symbols.gmt; c6.all.v6.1.symbols.gmt; c7.all.v6.1.symbols.gmt) was computed using standard settings, collapsing probe sets within genes using the Max_probe method and using 5000 permutations. For further analysis, we kept only gene sets that were significant in at least one of the up or down regulated genes in the two comparisons mentioned above at FDR < 5%.

